# The kinesin KIF3AC recycles endocytosed integrin to polarize adhesion formation towards the leading edge

**DOI:** 10.1101/2024.12.09.627580

**Authors:** Johnny A. Z. Rockenbach, Guilherme P. F. Nader, Susumu Antoku, Gregg G. Gundersen

**Author notes:** Correspondence should be addressed to G.G.G.

## Abstract

The recycling of integrin endocytosed during focal adhesion (FA) disassembly is critical for cell migration and contributes to the polarized formation of new FAs toward the leading edge. How this occurs is unclear. Here, we sought to identify the kinesin motor protein(s) that is involved in recycling endocytosed integrin back to the plasma membrane. We show that the kinesin-2 heterodimer, KIF3AC and the Rab11 adaptor protein RCP are required for FA reformation after the disassembly of FAs in mouse and human fibroblasts. In the absence of KIF3AC, integrin does not return to the cell surface after FA disassembly and is found in the Rab11 endocytic recycling compartment. Biochemical pulldowns revealed that KIF3C associated with β1 integrin in an RCP dependent fashion, but only after FA disassembly. KIF3AC knockdown inhibited cell migration, trafficking of RCP toward the leading edge, and polarized formation of FAs at the leading edge. These results show that KIF3AC promotes cell migration by recycling integrin so that it generates new FAs in a polarized fashion.

**Summary:** The study reveals that the heterodimeric kinesin-2 motor KIF3AC and its adaptor RCP are crucial for polarized formation of focal adhesions at the front of migrating fibroblasts. KIF3AC and RCP associate with intracellularly recycling integrin to promote its return to the cell surface after its endocytosis from disassembled focal adhesions.

## Introduction

Cell migration is an integral component of development, wound healing, immune cell responses and cancer metastasis (Luster et al., 2005; Scarpa and Mayor, 2016; Fanfone et al., 2022). Given the significance of cell migration for these processes, there has been substantial effort to understand the cellular processes that contribute to migration and how they are optimized for migration in different environments.

The adhesion of the cell to the substratum is one of the key steps in cell migration (Mitchison and Cramer, 1996; Ridley et al., 2003; Conway and Jacquemet, 2019). In both 2D and 3D environments, integrin-based adhesions are critical for migration, although there are integrin-independent modes of migration (Petrie et al., 2009; De Franceschi et al., 2015; Yamada and Sixt, 2019; Pawluchin and Galic, 2022). In response to extracellular matrix ligands, integrins cluster into small focal complexes that mature into larger, force bearing focal adhesions (FAs). These adhesions form in the front of the cell and detach in a stochastic manner behind the leading edge so that only a few survive to reach the cell rear (Smilenov et al., 1999; Möhl et al., 2012; Stehbens et al., 2014).

Disassembly of adhesions is just as important as assembly, as failure to disassemble keeps the cell stuck in place. A prominent mode of FA disassembly is microtubule-stimulated, clathrin-dependent endocytosis of integrins from adhesions (Ezratty et al., 2005, 2009; Chao and Kunz, 2009; Nader et al., 2016). The turnover of adhesions depends on their location: adhesions behind the leading edge and in the cell body turn over by microtubule-stimulated and clathrin-dependent endocytosis (Ezratty et al., 2005, 2009; Chao and Kunz, 2009; Nader et al., 2016), those in the cell rear detach by myosin II-dependent contraction (Vicente-Manzanares et al., 2009).

We know relatively little about the fate of integrins in adhesions in the cell rear. At least in some cases in 2D migration where there is strong adhesion, FAs in the cell rear along with a small amount of cytoplasm are ripped from the cell by contractile forces (Regen and Horwitz, 1992). This suggests that integrins in rear adhesions may not be reutilized in all cases.

In contrast, integrin derived from FAs disassembled by microtubule-stimulated, clathrin-mediated endocytosis is internalized and recycled in a Rab5-, Rab11- dependent process (Ezratty et al., 2005, 2009; Chao and Kunz, 2009; Nader et al., 2016). This ensures that the integrin is delivered to the endocytic recycling compartment (ERC) near the nucleus and then returned to the plasma membrane. The endocytosed integrin remains in an active conformational state during recycling, so that upon return to the cell surface, it is primed to reform adhesions (Nader et al., 2016). Consistent with this “integrin adhesion memory”, adhesions reformed from endocytosed integrin are polarized toward the leading edge of cells, but only when the endocytosed integrin is maintained in the active state (Nader et al., 2016). Sites of integrin exocytosis are primarily distributed toward the cell periphery adjacent to forming adhesions consistent with a close coupling between integrin arrival at the plasma membrane and incorporation into adhesions (Huet-Calderwood et al., 2017).

The polarized reformation of adhesions from endocytosed integrin near the leading edge has been proposed to contribute to the directional mobility of cells (Nader et al., 2016). For this to be the case, there must be a means to deliver the recycled integrin in close proximity to the leading edge. Most long-distance vesicle trafficking, including that of endocytosed proteins, occurs on microtubules and is mediated by microtubule motors including kinesin-1, -2 and -3 and cytoplasmic dynein (Hirokawa et al., 2009). Several of the kinesin motors selectively recognize post-translationally modified microtubules (Liao and Gundersen, 1998; Lin et al., 2002; Hammond et al., 2010; Li and Gundersen, 2008; Sirajuddin et al., 2014; McKenney et al., 2016), which themselves are polarized toward the leading edge (Gundersen and Bulinski, 1988; Li and Gundersen, 2008; Doyle et al., 2009). These stabilized and marked MTs span the distance from the centrosome to the leading edge in 2D and 3D migrating cells and so could selectively deliver cargoes, including endocytosed integrin to the leading edge. Indeed, there is evidence that transferrin receptors are recycled on stabilized and post-translationally modified microtubules (Lin et al., 2002).

What has been missing from the picture is how recycled integrins are trafficked internally in the cell toward the leading edge. Kinesins are logical candidates to serve this function and kinesins have been implicated in cell migration and in some cases integrin trafficking. The mitotic centromere-associated kinesin MCAK/KIF2C affects FA assembly and disassembly after nocodazole washout through an unknown mechanism (Moon et al., 2021). KIF15 regulates integrin internalization by delivering Dab2 to the plasma membrane and promoting FA disassembly (Eskova et al., 2014; He et al., 2022). KIF1C plays a role in detachment of adhesions toward the cell rear by delivering integrin to and maintaining the cell tail (Theisen et al., 2012). In none of these studies has a kinesin been identified that associates with endocytically-derived integrin and contributes to formation of FA near the leading edge.

Here, we identify the heterodimeric kinesin-2, KIF3AC as a kinesin motor involved in the recycling of integrin endocytosed after FA disassembly. We show that KIF3AC is specifically associated with endocytosed integrin through the Rab11 adaptor, Rab coupling protein (RCP) and both proteins are required for the reformation of adhesions after microtubule-induced FA disassembly. Consistent with its role in integrin recycling and FA reformation, we find that kinesin KIF3AC is required for cell migration into an in vitro wound, for the polarized trafficking of RCP endosomes, and to polarize adhesion formation at the front of migrating cells.

## Results

### The kinesin-2 motor KIF3AC is required for FA reassembly after FA disassembly

Kinesin vesicle motors (kinesin-1, -2, and -3) are candidates for recycling endocytosed integrin from the Rab11 recycling compartment (Hirokawa et al., 2009; Lin et al., 2002). We focused on kinesin-2, a heterodimeric kinesin composed of KIF3A and either KIF3B or KIF3C subunits (Yamazaki et al., 1995; Muresan et al., 1998; Marszalek and Goldstein, 2000), because it participates in Rab11 endocytic recycling of transferrin receptor and interacts with the Rab11 adaptor, Rab11-FIP5 (Schonteich et al., 2008a). KIF3A has also been shown to contribute to cell migration (Gao et al., 2020).

As an initial test of whether kinesin-2 might affect integrin adhesions, we knocked down the obligate subunit of kinesin-2 heterodimer, KIF3A in NIH3T3 fibroblasts and examined the cells for FAs by immunofluorescence of FA proteins (e.g., vinculin or paxillin). Knockdown of KIF3A with two distinct siRNAs reduced KIF3A protein level (Figure S1A) and the number of FAs detected by paxillin staining in steady state cells (Figure 1A). Both large and small adhesions were reduced in number in KIF3A depleted cells, but there was only a statistically significant reduction in size for one of the two KIF3A siRNAs (Figure 1B, C).

**Figure 1:**
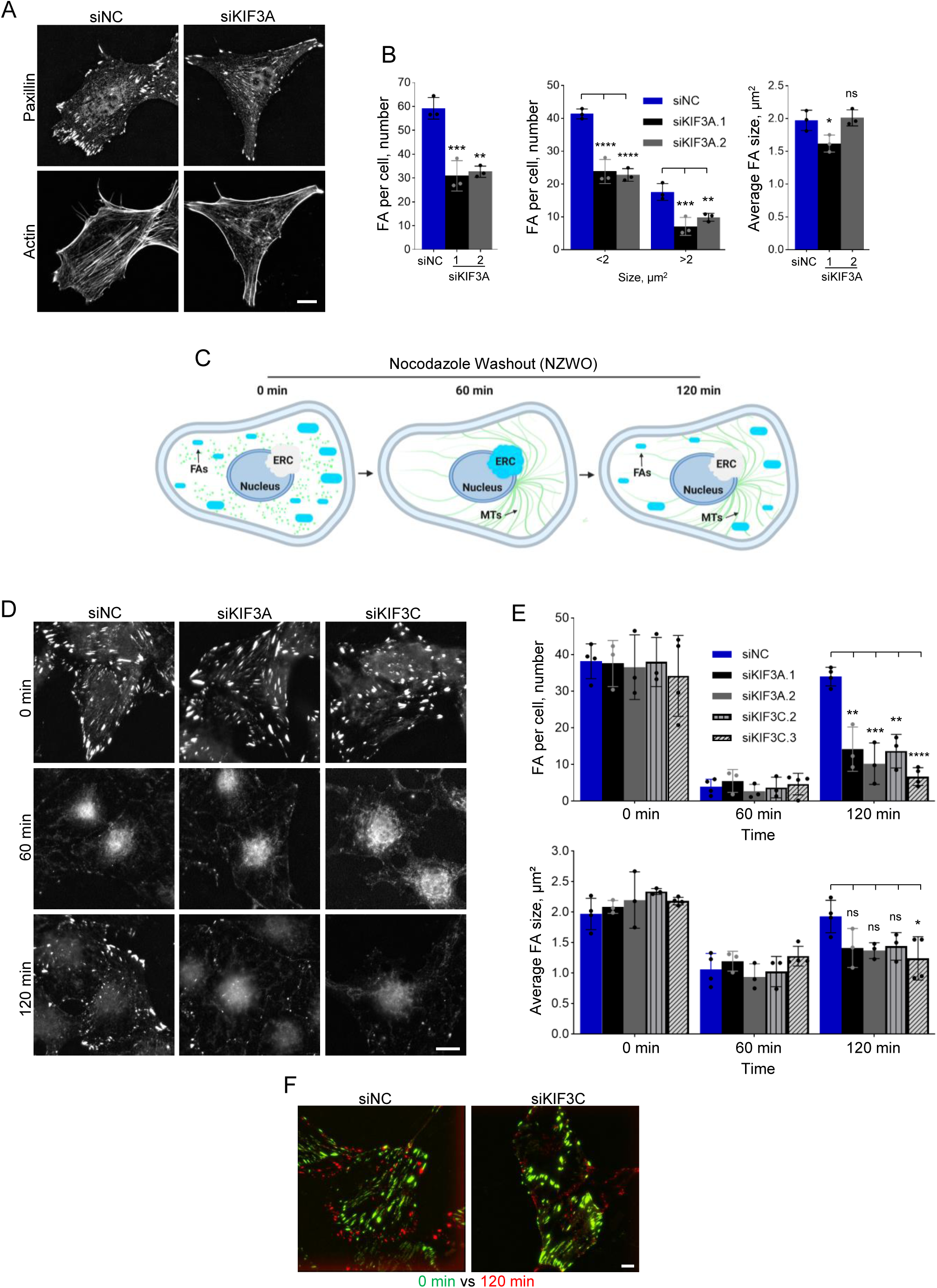
KIF3AC knockdown alters FA number and reassembly from endocytosed integrin. **A)** Representative images of steady state NIH3T3 fibroblasts transfected with noncoding siRNA (siNC) or siRNA for KIF3A (siKIF3A) and stained for FAs with paxillin antibody and filamentous actin with phalloidin. Scale bar, 15 µm. **B)** Quantification of FA number (left), categorized size (middle) and average size (right) for cells treated as in A. Error bars, mean ± SD; p values: *, < 0.05; **, < 0.01; ***, <0.005; ****, 0.001; NS, not significant. The histogram bars represent the standard deviation (SD), with each dot indicating the mean of an individual experiment. **C)** Schematic of nocodazole washout assay. Created with BioRender.com. **D)** Representative images from a nocodazole washout assay to assess microtubule-induced FA disassembly and reassembly. Nocodazole-treated NIH3T3 fibroblasts, previously depleted of the indicated proteins by siRNAs, were fixed at 0, 60 and 120 min after nocodazole washout and stained for paxillin and filamentous actin as in A. Scale bar, 15 µm. **E)** Quantification of FA number and size from cells treated as in D. Two different siRNAs were used for both KIF3A and KIF3C. Data in B and E represent 30 cells per condition from 4 independent experiments. The histogram bars bars represent the SD, with each dot indicating the mean of an individual experiment. **F)** The panels show overlaid frames of total internal reflection fluorescence movies of paxillin-EGFP expressing NIH3T3 fibroblasts at 0 min after nocodazole treatment (in green), and 120 min after nocodazole washout (in red). Cells were treated with the indicated siRNAs. For the complete movie, see Supplementary Movie 1.

To test whether KIF3A affected FA assembly, we quantified the number and size of adhesions following lysophosphatidic acid (LPA) stimulation of serum-starved fibroblasts, a classic assay of contractility-induced FA formation (Ridley and Hall, 1992) (Figure S1D). At either 15 min or 90 min after LPA-stimulation, the average number or size of paxillin-labeled adhesions per cell was unaffected by KIF3A knockdown (Figure S1D, E). This suggests that KIF3A does not participate in the contractile assembly of adhesions.

To test whether KIF3A is involved in FA disassembly or the subsequent endocytic recycling, we used the nocodazole washout assay (Ezratty et al., 2005, 2009; Yeo et al., 2006; Chao and Kunz, 2009; Nader et al., 2016). In this assay, MT regrowth after nocodazole removal triggers the synchronous disassembly of FAs by an endocytic process so that by 60 min almost all FAs disassemble. Following disassembly, the endocytosed integrin synchronously recycles to the plasma membrane where it contributes to the de novo formation of new FAs (see diagram, Figure 1C). KIF3A knockdown by two distinct siRNAs (Figure S1A) did not interfere with the disassembly of FAs induced by microtubule regrowth (t=60 min) but strongly prevented the reformation of FAs 120 min after microtubule regrowth, when controls assembled new adhesions (Figure 1D, E). FA number, but not size, was affected (Figure 1E). Kinesin-2 is an obligate heterodimer of KIF3A and either KIF3B or KIF3C (Marszalek and Goldstein, 2000). To determine which heterodimer of kinesin-2 is involved in FA reassembly, we knocked down KIF3B and KIF3C and confirmed reduced protein levels by western blot (Figure S1B, C). Knockdown of KIF3B did not affect the disassembly of FAs triggered by microtubule regrowth (t=60 min) or the subsequent reassembly of adhesions (t=120) (Figure S1F-H). Knockdown of KIF3C also did not affect FA disassembly, but like KIF3A knockdown, it significantly inhibited the reassembly of FAs 120 min post-nocodazole washout (Figure 1G, H). As with KIF3A, knockdown, KIF3C knockdown affected FA number, but not size (Figure 1E). None of the KIF3 knockdowns affected MT regrowth (Figure S1F).

To follow this process dynamically, we monitored paxillin-GFP as a marker of FAs and prepared movies of NIH3T3 fibroblasts after washing out nocodazole. These movies show FAs disassembling after nocodazole washout in both controls treated with noncoding siRNA (siNC) and in cells treated with siKIF3C (Supplementary Movie 1). Overlay of the first frame (0 min nocodazole washout) and last frame (120 min post-nocodazole washout) from these movies clearly shows FAs returning after nocodazole washout in the control cells, but not in the siKIF3C treated cells (Figure 1F). As reported previously, the newly reformed FAs after nocodazole washout did not return to the same location as the original FAs before nocodazole washout (Nader et al., 2016).

To investigate whether KIF3AC was also involved in FA reassembly in other cells, we conducted similar nocodazole washout assays with primary human dermal fibroblasts (HDF). HDFs treated with noncoding shRNA (shNC) (Figure S2A) behaved similarly to NIH3T3 fibroblasts in the nocodazole washout assay: FAs disassembled by 45-60 min after nocodazole washout and reassembled 120 min after nocodazole washout (Figure S2B). Similar to NIH3T3 fibroblasts, HDFs treated with KIF3C shRNA disassembled their FAs after nocodazole washout (t=45) but failed to reassemble FAs (t=120) (Figure S2B, C). As in NIH3T3 fibroblasts, the number of FAs reformed was reduced in KIF3C-depleted HDFs without significantly affecting their size (Figure S2C). Taken together, these results demonstrate that the kinesin KIF3AC plays a role in FA reassembly following endocytic disassembly of FAs, suggesting that it may be involved in the trafficking of the endocytosed integrins after FA disassembly.

### KIF3A and KIF3C tails inhibit FA reassembly

The head of kinesin-2 binds to microtubules whereas the tail binds to cargos (Li et al., 2014). To test whether acute inhibition of KIF3AC also affects FA reassembly, we microinjected a tail construct lacking the head domain of KIF3A (KIF3A-tail-EGFP) and performed the nocodazole washout assay. Neither KIF3A-tail-EGFP nor EGFP microinjection affected FA disassembly induced by nocodazole washout (t=60 min, Figure 2A, B). In contrast, microinjected KIF3A-tail-EGFP impaired FA reassembly when compared to EGFP-microinjected cells (t=120 min, Figure 2A, B). We observed similar inhibition of FA reassembly, but not disassembly after nocodazole washout in cells expressing dominant negative KIF3C-tail by transfection (Figure 2C, D). These results show that acutely interfering with KIF3AC with dominant negative constructs inhibits FA reassembly in a similar fashion as longer-term depletion of the protein by knockdown and suggest that the cargo binding domain of KIF3AC is involved.

**Figure 2:**
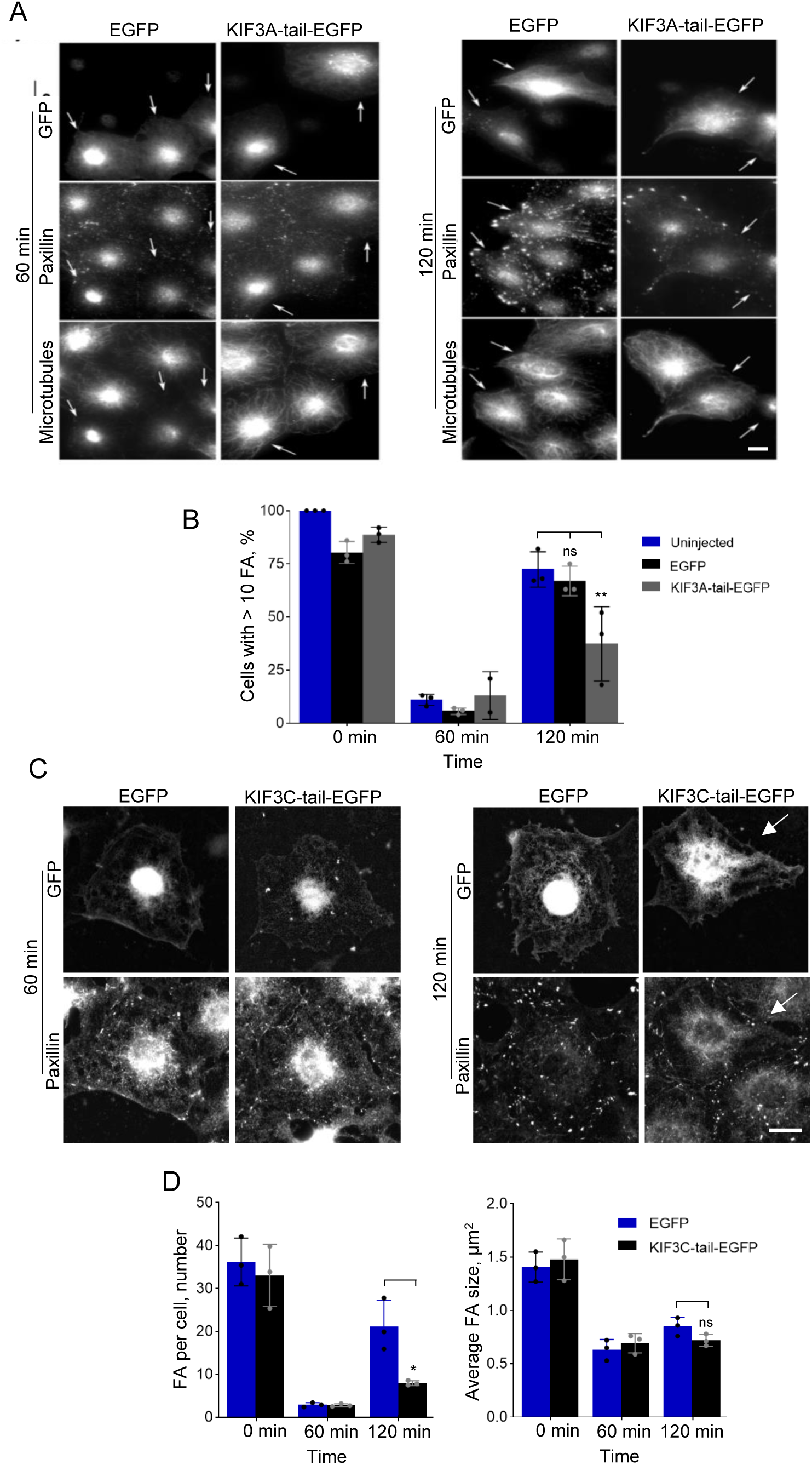
Dominant negative KIF3AC tail constructs block FA reassembly. **A)** Representative images of NIH3T3 fibroblasts microinjected with either EGFP or KIF3A-tail-EGFP and fixed at the indicted times after nocodazole washout. Cells were stained for paxillin, tubulin and GFP. Scale bar, 15µm. **B)** Quantification of microinjected cells exhibiting > 10 FAs at the indicated times after nocodazole. Cells expressed the indicated constructs as in A. At least 35 cells were quantified for each time point in 3 different experiments. The histogram bars represent the SD, with each dot indicating the mean of an individual experiment. **C)** Representative images of NIH3T3 fibroblasts transfected with either EGFP or KIF3C-tail-EGFP and fixed at the indicted times after nocodazole washout. Cells were stained for paxillin and GFP. Scale bar 15µm. **D)** Quantification of FA number and size for the transfected cells at the indicated times after nocodazole washout. Thirty cells were quantified at each time point from 3 independent experiments. The histogram bars represent the SD, with each dot indicating the mean of an individual experiment.

### KIF3A is required for integrin trafficking from the endocytic recycling compartment (ERC) to the cell surface

FA reassembly from endocytosed integrin coincides with and requires the recycling of endocytosed α5β1 integrin to the cell surface (Nader et al., 2016). To test whether KIF3AC is required for the recycling of endocytosed integrin to the cell surface, we examined cell surface levels of endogenous α5 integrin by flow cytometry of cells fixed during FA disassembly and reassembly triggered by microtubule regrowth (Figure 1C). KIF3A-depleted cells reduced their surface α5 integrin levels to the same extent as control cells after FA disassembly (60 min post-nocodazole washout) but failed to return surface α5 integrin levels at the time of FA reassembly (120 min post-nocodazole washout) (Figure 3A). This result supports the conclusion that KIF3A is required to recycle the integrin endocytosed from FAs.

**Figure 3:**
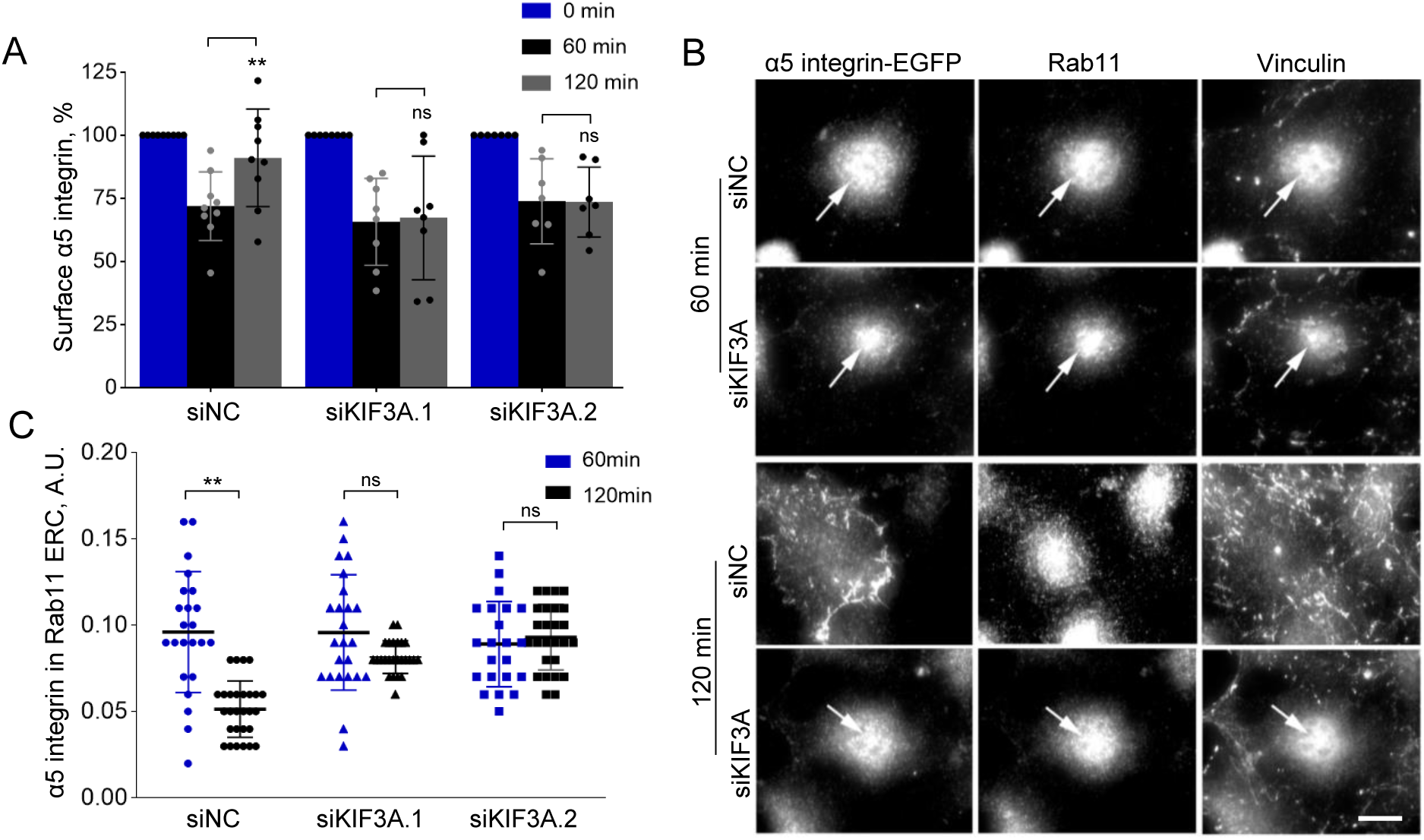
KIF3AC is required to recycle integrin to the plasm membrane after FA disassembly. **A)** Levels of surface α5 integrin during nocodazole washout in NIH3T3 fibroblasts treated with either noncoding or KIF3A siRNAs. Surface levels were determined by FACS and are plotted relative to the levels at 0 min of nocodazole washout. For each condition, 10,000 cells were analyzed in at least 7 independent experiments. **B)** Representative images showing α5-integrin- EGFP colocalization with the Rab11 compartment at 60 and 120 min after nocodazole washout. Scale bar 15µm. **C)** Quantification of α5 integrin-EGFP in the Rab11 compartment from images as in A. At least 22 cells per condition were quantified from 3 independent experiments.

To determine where the integrin resided in KIF3AC-depleted cells, we localized expressed α5 integrin-EGFP after FA disassembly. As reported previously (Nader et al., 2016), in controls, α5 integrin-EGFP transiently accumulated in the pericentriolar ERC at 60 min after nocodazole washout, but at later times (120 min) when FAs had reformed, levels of α5 integrin-EGFP in the ERC were reduced (Figure 3B, C). In contrast, in KIF3A-depleted cells, α5 integrin-EGFP localized in the ERC at both the 60 min and 120 min time points (Figure 3B, C).

Consistent with previous results showing certain FA components travel with endocytosed integrin (Nader et al., 2016), we detected vinculin in the ERC with the endocytosed integrin during recycling. In controls, vinculin was observed at the ERC at 60 min after FA disassembly but was present in FAs after they reassembled at 120 min (Figure 3B). In contrast, vinculin was observed in the ERC at both the 60 and 120 min time points after FA disassembly in KIF3A depleted cells (Figure 3B, C). These results indicate that in the absence of KIF3A, integrin traffics to but does not move out of the ERC to the cell surface as in control cells.

### KIF3A colocalizes with integrin in the Rab11 ERC after FA disassembly

To determine how directly KIF3AC might be affecting integrin recycling, we first localized endogenous KIF3A after FA disassembly. KIF3A colocalized with Rab11 in the ERC 60 min after nocodazole washout (Figure 4A). At 120 min post-nocodazole washout when FAs had reassembled, the level of KIF3A colocalizing with Rab11 in the ERC dropped (Figure 4A, B). KIF3A did not colocalize with reformed FAs, suggesting it was associated only during intracellular trafficking (Figure 4A, B). To determine if this KIF3A localization coincided with recycling integrin, we monitored integrin distribution by expressing α5-integrin-EGFP. Both α5-integrin-EGFP and KIF3A were detected in the ERC 60 post-nocodazole washout and both showed decreased ERC localization 120 min post-nocodazole washout when FAs had reformed (Figure 4C). Together, these results show that KIF3A localization in the ERC coincides with integrin derived from FA disassembly.

**Figure 4:**
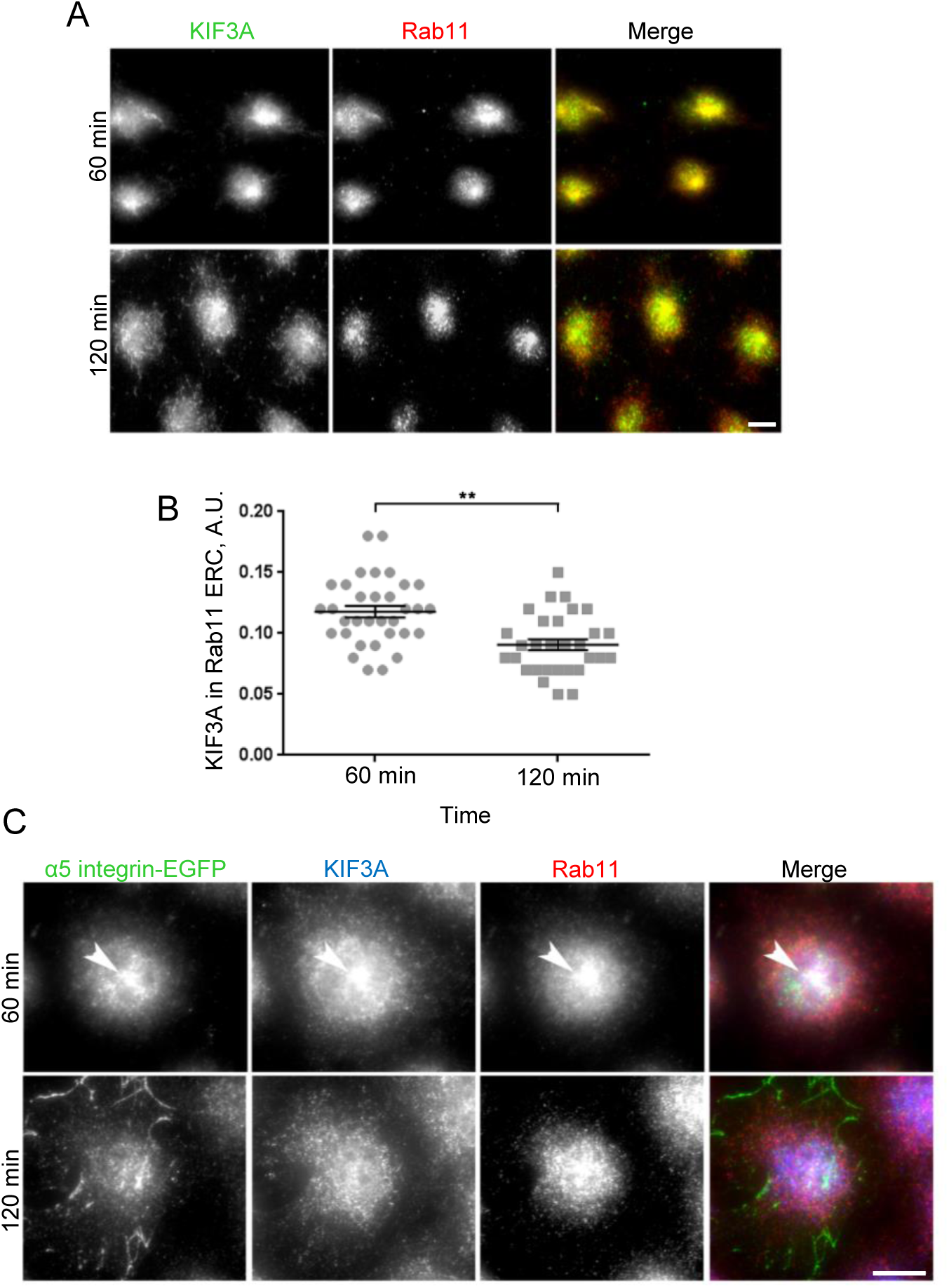
KIF3A localizes in the Rab11 ERC following FA disassembly. **A)** Representative images of endogenous KIF3A colocalization with the Rab11 ERC 60 and 120 min after nocodazole washout in NIH3T3 fibroblasts. Scale bar, 15 µm. **B)** Quantification of KIF3A in the Rab11 compartment from images stained as in A. At least 30 cells in each experiment were quantified from 3 independent experiments. **C)** Colocalization of α5 integrin-EGFP and KIF3A in the Rab11 compartment at 60 and 120 minutes after nocodazole washout in NIH3T3 cells. Arrowheads indicate colocalization of α5 integrin-EGFP and KIF3A with Rab11 at 60 min. Scale bar, 15 µm.

### KIF3AC biochemically associates with Integrin and talin only during endocytic recycling

The requirement of KIF3AC for integrin recycling and its localization with integrin in the ERC only during recycling support the hypothesis that KIF3AC acts as a molecular motor for integrin recycling after FA disassembly. To test this hypothesis further, we determined whether KIF3AC biochemically associated with recycling integrin after FA disassembly. We infected either HEK- 293T cells with viruses encoding KIF3C-GST or GST as a control together with β1-integrin-myc and performed GST pull-downs. Western blots of the GST-pull downs in HEK-293T cells revealed that β1-integrin-myc specifically associated with KIF3C-GST, but only after nocodazole washout to trigger FA disassembly (Figure 5A). Consistent with previous results showing that certain FA proteins associate with endocytosed integrin (Nader et al., 2016), endogenous talin was also present in the KIF3A-GST pull downs (Figure 5A). Moreover, in NIH3T3 fibroblasts expressing KIF3C-GST, endogenous β1-integrin and talin also specifically associated with KIF3C after nocodazole washout (Figure 5B). Association between KIF3C-GST and β1-integrin was first detected at 15-30 min after nocodazole washout but increased at 60 min coincident with FA disassembly and integrin localization in the ERC (Figure 5B). These results establish that KIF3AC associates with integrin in a manner that depends on the disassembly of FAs and strongly support the idea that KIF3AC is a motor protein for recycling endocytosed integrin.

**Figure 5:**
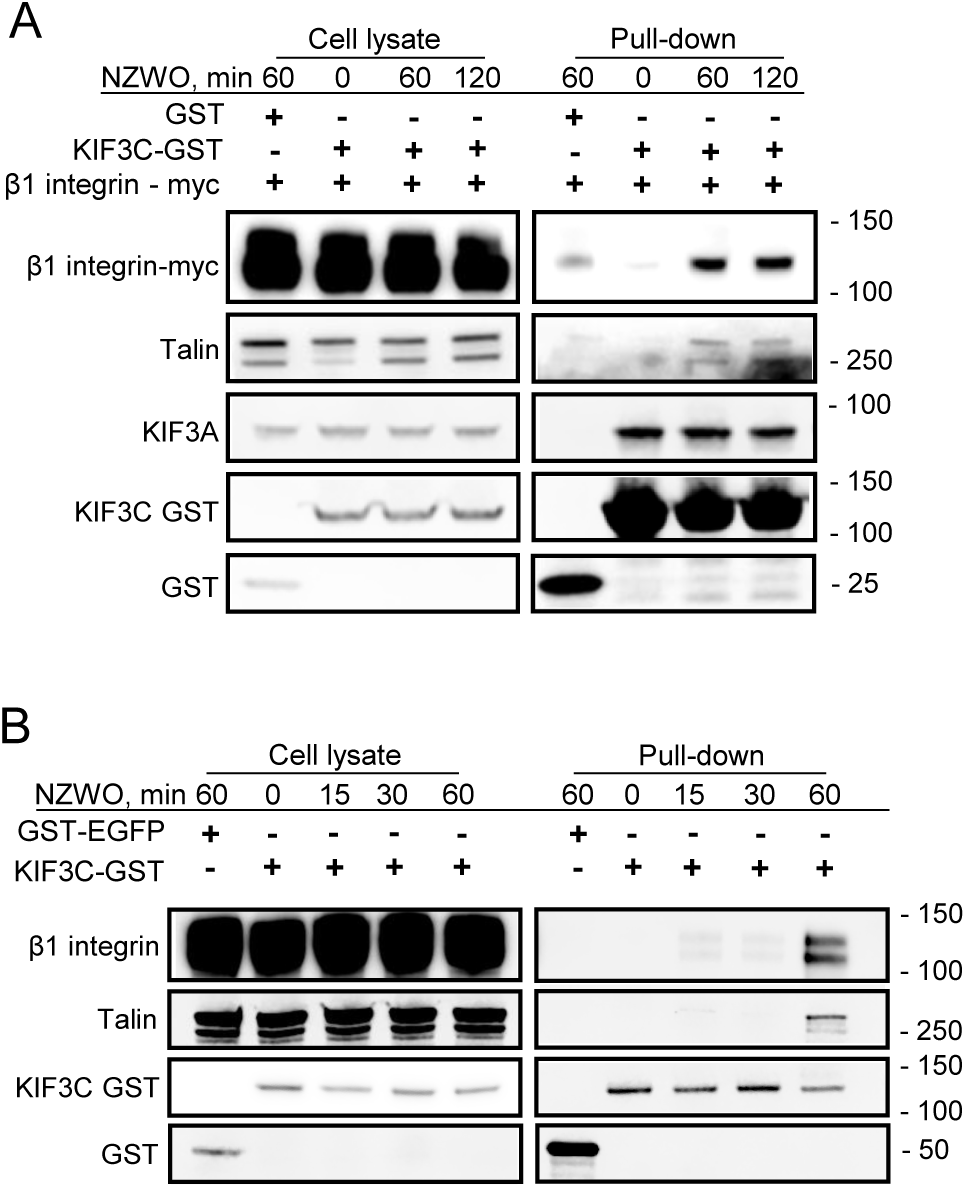
KIF3C specifically associates with β1-integrin and talin after FA disassembly. **A)** Western blots of cell lysates and GST pulldowns from HEK293T cells expressing β1-integrin-myc and GST or KIF3C-GST. Samples were prepared at the indicated times after nocodazole washout (NZWO). **B)** Western blots of cell lysates and GST pulldowns from NIH3T3 cells expressing either GST-EGFP or KIF3C-GST. Western blots were probed with the antibodies indicated on the left. Samples were prepared at the indicated times after nocodazole washout (NZWO). Molecular weight markers are on the right in kDa.

### Rab coupling protein (RCP/Rab11-FIP1) is required for the association between integrin, talin and KIF3AC

Most vesicular motor proteins, including kinesin-2, associate with their cargoes through adaptor proteins (Gilbert et al., 2018). We tested three kinesin-2 adaptor proteins: KAP3 (Garbouchian et al., 2022) and two Rab11 binding adaptors, Rip11/Rab11-FIP5 (Schonteich et al., 2008) and RCP/Rab11-FIP1 (Caswell et al., 2008), to see if one of them affected FA reassembly after nocodazole washout. KAP3 or Rip11/Rab11-FIP5 knockdown both had no effect on either FA disassembly or reassembly after nocodazole washout (Figure S3A, B, D, E). Knockdown was confirmed by western blotting (Figure S3C, F).

We next tested RCP/Rab11-FIP1, which was previously shown to be involved in integrin recycling (Caswell et al., 2008). We find that the knockdown of RCP (Figure S3G) blocked FA reassembly after nocodazole washout (t=120 min), while having no effect on FA disassembly (t=60 min) (Figure 6A, B). Live cell movies with NIH3T3 cells expressing paxillin-EGFP also showed that in single cells, RCP knockdown did not prevent FA disassembly, but blocked reformation of FAs when compared to control cells (Figure 6C, Supplementary movie 2). Next, we tested whether

**Figure 6:**
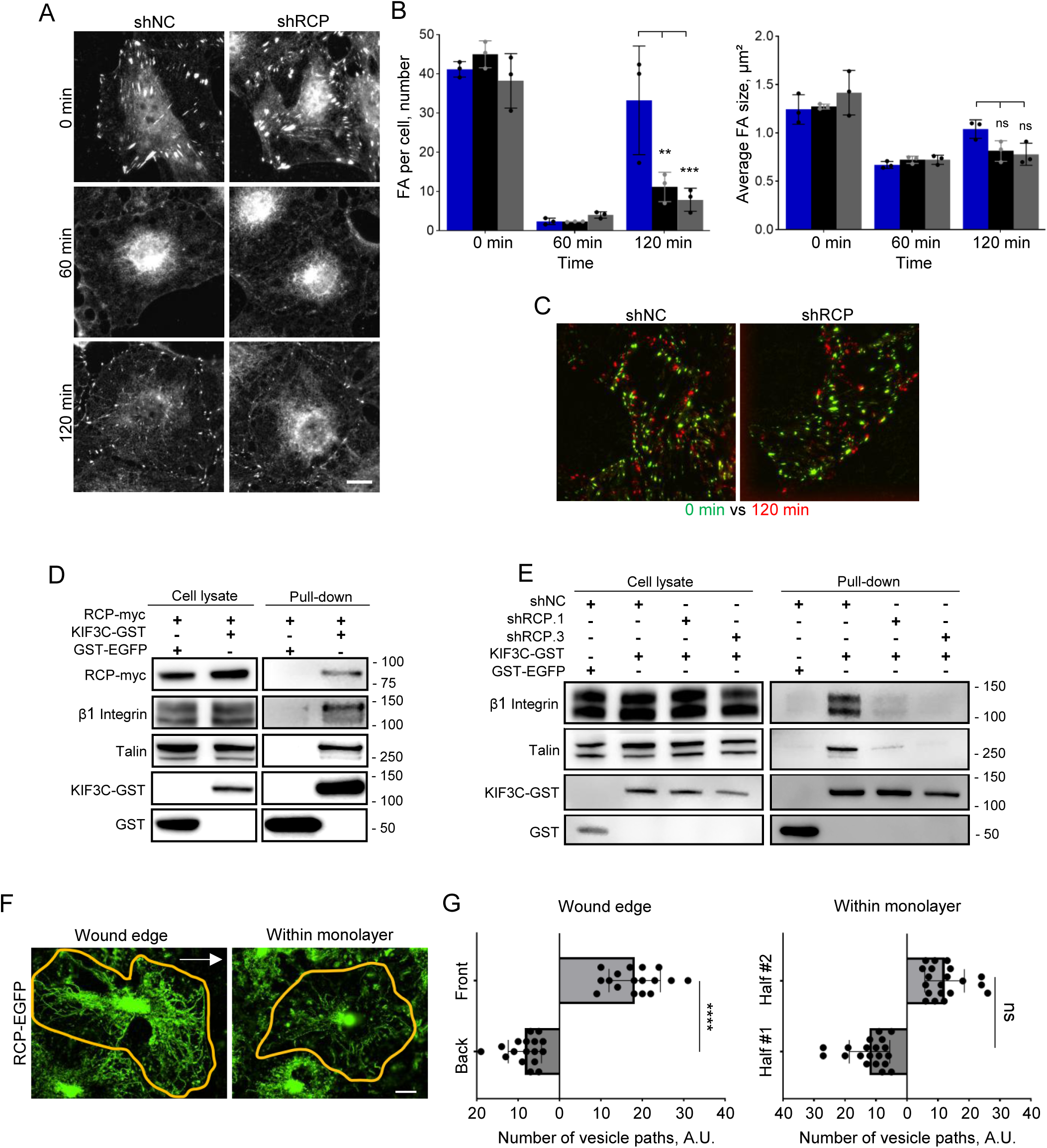
RCP/Rab11-FIP1 is required for FA reassembly and the association between β1-integrin and talin with KIF3C. **A)** Representative images of FAs stained with paxillin antibody in NIH3T3 fibroblasts at the indicated times after nocodazole washout. Cells were transfected with the indicated shRNAs. Scale bar, 15 µm. **B)** Quantification of paxillin-stained FA number and size in shNC- or shRCP-transfected NIH3T3 fibroblasts fixed at the indicated times after nocodazole washout. Data are from 90 cells per time point from three independent experiments. The histogram bars represent the SD, with each dot indicating the mean of an individual experiment. **C)** The panels show overlaid frames of total internal reflection fluorescence movies of paxillin-EGFP expressing NIH3T3 fibroblasts at 0 min after nocodazole treatment (in green), and 120 min after nocodazole washout (in red). Cells were treated with the indicated shRNAs. For the complete movie, see Supplementary Movie 2. **D)** Western blots of cell lysates and GST pulldowns from NIH3T3 cells expressing RCP-myc and either KIF3C-GST or GST-EGFP. Samples were prepared 60 minutes after nocodazole washout (NZWO). **E)** Western blots of cell lysates and GST pulldowns from NIH3T3 cells expressing the indicated shRNAs and either KIF3C-GST or GST-EGFP. Samples were prepared 60 minutes after nocodazole washout (NZWO). Western blots were probed with the antibodies indicated on the left. Molecular weight markers are on the right in kDa. **F)** Maximum projection overlays representing 5 min of movies prepared from RCP-GFP expressing NIH3T3 fibroblasts 60 min after nocodazole washout. The projections reveal RCP vesicle paths in polarized cells at the wound edge (arrow denote direction to wound edge) and nonpolarized cells within the monolayer. Cell boarders are drawn in yellow. Projections are from Supplementary Movie 3. **G)** Quantification of the vesicle path directions for cells as in F. At least 15 cells were analyzed in each condition from 4 independent experiments. Cells were divided into equal halves. In wound edge cells, the two halves represent the front and back of the cell; in cells within the monolayer, the halves were arbitrarily draw and represented as half #1 and half #2.

RCP associated with KIF3C after FA disassembly (t=60 min). For this, we expressed both RCP-myc and KIF3C-GST and performed pulldowns from samples prepared during nocodazole washout. We find that RCP-myc specifically associated with KIF3C-GST after FA disassembly (t=60 min) (Figure 6D). Endogenous β1-integrin and talin were also detected in the KIF3C-GST pull-downs (Figure 6D), consistent with RCP acting as an adaptor between integrin and KIF3AC.

To directly test if RCP acted in a manner consistent with it being a motor adaptor for integrin and KIF3AC, we conducted KIF3C-GST pull downs in NIH3T3 fibroblasts expressing two different shRNAs for RCP, both of which reduced endogenous RCP levels (Figure S3G). Expression of either RCP shRNAs substantially decreased the association between KIF3C-GST and both integrin and talin detected after FA disassembly (Figure 6E). Thus, RCP is a specific adaptor for coupling KIF3AC to endocytosed integrin.

### RCP vesicles move in a polarized fashion toward the leading edge

Given our data showing that RCP associates with β1 integrin and KIF3C (Figure 6D) during nocodazole washout, we utilized RCP-EGFP to monitor vesicle movements during this process. In this approach we are using RCP as a surrogate as detecting vesicles with integrin or KIF3C in NIH3T3 fibroblasts was not feasible.

During cell migration, focal adhesions form at the front of the cell to lead and propel migration (Lauffenburger and Horwitz, 1996; Gardel et al., 2010). In a wounded monolayer, integrin delivery is expected to be polarized towards the leading edge. We hypothesized that RCP trafficking would also be polarized towards the wound edge. To test this hypothesis, we expressed RCP-EGFP in NIH3T3 fibroblasts during the nocodazole washout assay and imaged different cells every 400 ms for 5 minutes, ranging from 60 to 120 minutes after nocodazole washout. We prepared movies of RCP-EGFP trafficking in cells at the wound edge to represent polarized cells and compared these to cells that were within the monolayer to represent nonpolarized cells (Figure 6F, G, Supplementary movie 3). Our results show that while non-polarized cells within the monolayer exhibited non-polarized trafficking of RCP vesicles, cells at the wound edge demonstrated a clear polarization of vesicle paths towards the leading edge.

### KIF3AC is critical for polarized reassembly of FAs in migrating cells

KIF3A has been implicated in cell migration of MDCK and glioma cells (Boehlke et al., 2013; Gao et al., 2020). We confirmed that KIF3A or KIF3C knockdown inhibited the migration of the NIH3T3 fibroblasts used in this study (Figure 7A, B). The overall number of FAs in the migrating NIH3T3 fibroblasts were unaffected by KIF3C knockdown, but the size of individual FAs was significantly smaller (Figure 7C, D). Moreover, FAs in the leading edge of KIF3C-depleted cells were substantially smaller than in controls (Figure 7E, F) and the overall polarization of FAs toward the leading edge was reduced (Figure 7G, H). These results indicate that KIF3AC integrin recycling contributes to the selective formation of FA in the leading edge, consistent with previous work that showed recycled active integrin contributed to the formation of FAs at the leading edge (Nader et al., 2016).

**Figure 7:**
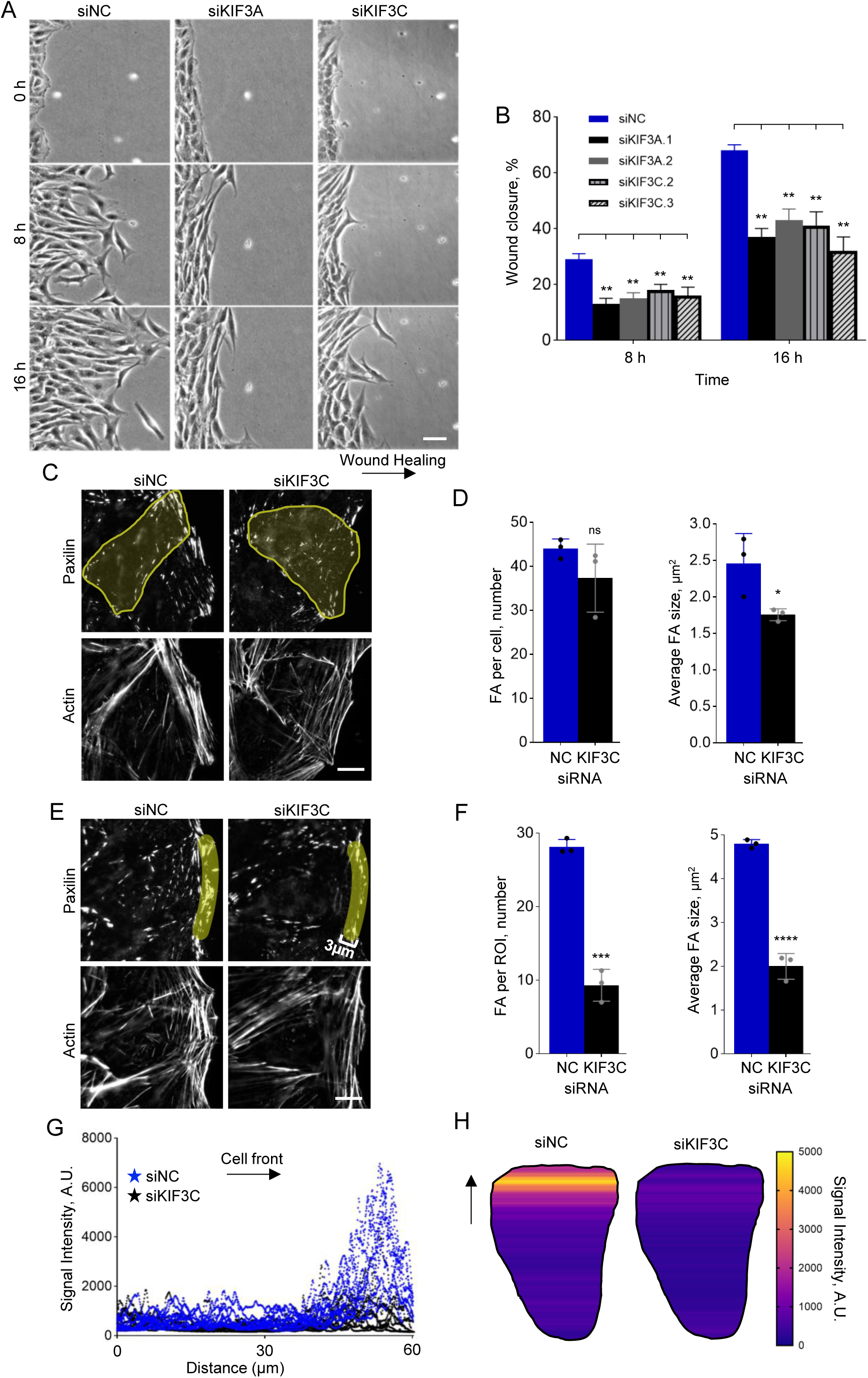
KIF3AC is critical for polarized reassembly of FAs in migrating cells. **A)** Wound healing assay of NIH3T3 fibroblasts treated with the indicated siRNAs. Scale bar 50µm. **B)** Quantification of wound healing as in A. Bars are SE from 3 independent experiments. **C)** Representative images of paxillin-stained FAs and filamentous actin in NIH3T3 fibroblasts migrating into an in vitro wound. Cells were treated with the indicated siRNAs. Scale bar, 15 µm. **D)** Quantification of FA number and size for cells treated as in B. Data are from 90 cells per condition from three independent experiments. The histogram bars represent the SD, with each dot indicating the mean of an individual experiment. **E)** Representative images of paxillin-stained FAs and filamentous actin in NIH3T3 fibroblasts migrating into an in vitro wound. Yellow area represents the 3 µm deep, leading edge region that was quantified. Cells were treated with the indicated siRNAs. Scale bar, 15 µm. **F)** Quantification of FA size and number at the cell leading edge (3µm), as shown in E. Thirty cells were quantified from 3 independent experiments. The histogram bars represent the SD, with each dot indicating the mean of an individual experiment. **G)** Line traces of relative signal intensity of FA distribution along the front-back cell axis in NIH3T3 fibroblasts migrating into an in vitro wound after treatment with the indicated siRNAs. Ten cells were quantified from 2 independent experiments. **H)** Heat map depicting the signal intensity of FA distribution from analysis shown in G. Arrow, direction of cell front. In G & H, A.U. is arbitrary units.

## Discussion

Integrin trafficking and FA dynamics are required for proper cell migration in many contexts. FA assembly and disassembly have been well-studied, and much is known about how they form and disassemble. Connecting these two processes is the recycling of endocytosed integrins to the plasma membrane where they can contribute to the formation of new adhesions (Caswell and Norman, 2008; Bridgewater et al., 2012; De Franceschi et al., 2015; Nader et al., 2016). However, there remains much we do not know about the intracellular recycling pathway for integrins, including the kinesin motor responsible for the return of endocytosed integrin to the plasma membrane (De Franceschi et al., 2015). Our results here show that the kinesin-2 motor KIF3AC is required for FA reformation after nocodazole washout and for the return of endocytosed integrin to the plasma membrane. KIF3AC colocalizes with endocytosed integrin in the Rab11 ERC and associates with endocytosed integrin after FA disassembly. The KIF3AC-integrin association requires the Rab11 adaptor RCP, a protein previously implicated in integrin recycling (Caswell et al., 2008). All of these data points to a model in which KIF3AC is a motor protein required for the transit of endocytosed integrin to the plasma membrane where it contributes to the reformation of FAs (Figure 8).

**Figure 8:**
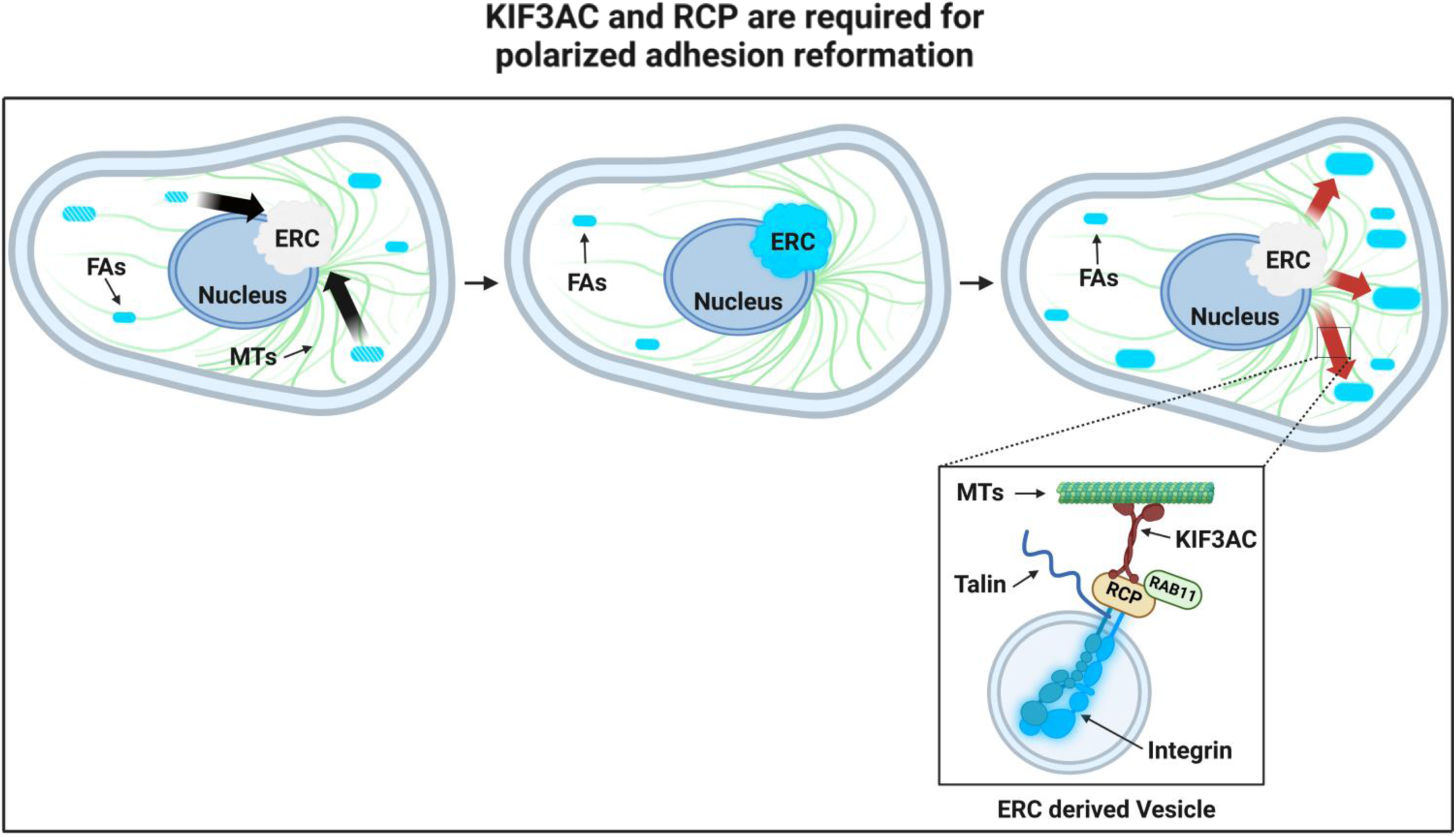
Schematic model for KIF3C associations with recycled integrin through RCP to polarize FA reformation at the leading edge. Integrins and associated proteins (e.g., talin) resident in the FA (e.g., talin) are endocytosed during FA disassembly and accumulate in the ERC before trafficking back to the plasma membrane by KIF3AC and RCP to polarize FA formation. Insert shows proposed relationship between endocytosed integrin and talin and the microtubule (MT) motor KIF3AC and its adaptor protein RCP. Created with BioRender.com.

Our results show that the association of KIF3C with integrin and associated FA proteins only occurs after FA disassembly (Figure 5A, B). These data points towards a switch in which the integrin in the FA cannot interact with KIF3AC, but upon disassembly of FAs, the endocytosed integrin and associated FA proteins become competent to associate with KIF3AC to allow intracellular trafficking. Rab11 is likely to be a key component of this switch. Rab11 is activated late in the recycling of endocytosed proteins (Wilcke et al., 2000; Ferro et al., 2021) and has been implicated in integrin recycling in the current study and numerous prior studies (Caswell and Norman, 2008; Ezratty et al., 2009; Nader et al., 2016). Rab11 works through Rab11 binding proteins (Wilcke et al., 2000; Eva et al., 2010; Wilson et al., 2023) and our data strongly suggest that for endocytosed integrin, Rab11 works through RCP. Importantly, we find that the behavior of RCP is consistent with it being the specific adaptor for endocytosed integrin traveling through the Rab11 pathway and that the other KIF3 adaptors Rab11-FIP5 and KAP3 do not seem to be required. This points to specificity at the level of the coupling protein, and it will be interesting to determine how RCP specificity for integrins is determined and whether it directly binds integrin tails or associates with integrin through integrin binding proteins, like talin.

There is substantial evidence that recycled integrins play a crucial role in formation of new FAs, particularly in the context of migrating cells (Caswell and Norman, 2008; De Franceschi et al., 2015; Moreno-Layseca et al., 2019). The present study extends this idea by showing that in the absence of KIF3AC and active integrin recycling, cells migrating into an in vitro wound have reduced FAs particularly at the leading edge and reduced rate of migration (Figures 7 and 8). These findings are consistent with those from a previous study indicating that in the absence of FAK activity to maintain endocytosed integrin in an active conformational state during trafficking, adhesions reform in a random fashion rather than polarized toward the leading edge (Nader et al., 2016). That KIF3AC is involved in this polarized reassembly of leading-edge FAs is also interesting given that its mobility on microtubules is one of the most strongly affected by post translationally detyrosinated tubulin (Sirajuddin et al., 2014). Detyrosinated tubulin accumulates in stabilized microtubules (Gundersen et al., 1984; Webster et al., 1987), and in migrating cells, these modified and stabilized microtubules are polarized toward the leading edge (Gundersen and Bulinski, 1988).

To further test the importance of recycled integrin in leading edge adhesion formation, it will be critical to identify sites of recycled integrin exocytosis relative to newly formed adhesions. Calderwood and colleagues showed convincingly that integrin exocytic sites were biased toward preexisting FAs, in support of directed delivery of trafficked integrin (Huet-Calderwood et al., 2017). However, their study did not identify the source of the exocytic integrin (biosynthetic vs endocytic), so it is still an open question where recycled integrin is delivered.

Our study raises many new questions about how integrin switches between its adhesion state and its intracellular trafficking state for recycling. From this study and an earlier one (Nader et al., 2016), we know that a number of focal adhesion proteins, such as talin, FAK, PIP1Kγi2 and vinculin travel with the recycling integrin and are detected with integrin in the ERC. Given talin’s interaction with the intracellular tail of β integrin subunits, it is possible that it mediates the connection to RCP. Kinesin-2 is a microtubule plus end-directed motor, so it is likely that endocytosed integrin needs to first associate with a minus end-directed motor, such as cytoplasmic dynein, to bring it to the Rab11 ERC. If this is the case, it is possible that there is an earlier switch in the association of the endocytosed integrin with microtubule motors and the KIF3AC-RCP switch we document here is temporally the second one in integrin’s intracellular travels. As far as we are aware, there are no studies of intracellular integrin trafficking by minus end motors, so this would be a fruitful avenue for future research.

## Material and Methods

### Antibodies

Antibodies used for Western Blot (WB) or Immunofluorescence (IF): β1 integrin (rabbit polyclonal, 1:100 dilution for IF, Abcam), pY118/31paxillin (rabbit polyclonal, 1:100 dilution for IF; Thermo Fisher Scientific), GFP (chicken polyclonal, 1:100 dilution for IF, EMD Millipore), KIF3A (mouse monoclonal, 1:1000 dilution for WB, BD Bioscience or rabbit polyclonal, 1:100 dilution for IF, MilliporeSigma), KIF3C (rabbit polyclonal, 1:1000 dilution for WB, MilliporeSigma), KIF3B (rabbit polyclonal, 1:1000 dilution for WB, MilliporeSigma), KAP3 (mouse monoclonal, 1:1000 dilution for WB, BD Bioscience), Rab11 (rabbit polyclonal; 1:200; BD Biosciences), vinculin (mouse monoclonal, 1:100 dilution for IF; MilliporeSigma), Paxillin (mouse, monoclonal, 1:200 dilution for IF, BD Bioscience), Rab11FIP5 (rabbit polyclonal, 1:200 dilution for IF, Thermo Fisher Scientific), Rab11FIP1 (rabbit polyclonal, 1:200 dilution for IF, Cell signaling), GST (mouse monoclonal, 1:2000 dilution for WB, Santa Cruz Biotech), Myc (rabbit polyclonal, 1:2000 dilution for WB, Cell Signaling), tyrosinated alpha-tubulin (rat monoclonal YL1/2, 1/10 dilution of culture supernatant for IF and 1:50,000 dilution for WB). Secondary antibodies for IF were from Thermo Fisher Scientific (donkey anti-rabbit and donkey anti-mouse, Alexa Fluor 488, 568, 648) and Abcam (goat anti-mouse DyLight 488). Secondary antibodies for WB were from Santa Cruz Biotech (goat anti-mouse and goat anti-rabbit, Horseradish peroxidase (HRP) conjugated) and LI-COR (goat anti-rabbit and goat anti-mouse, IRDye 800CW, IRDye 680RD).

### Plasmids, primers, shRNA and siRNA

KIF3A-tail-EGFP was a generous gift from Dr. Geri Kreitzer (CUNY, New York). pEF1a-GST-N4, pMSCV-GST-P-N4, and pMSCV-GST-6P-4 EGFP were previously described (Antoku et al., 2019). pEF1a-MYC-N4 was derived by inserting MYC tag sequence in the downstream of EF1a promotor in pEF1a vector (Antoku et al., 2019). pMSCV-puro-3xMYC-N4 was derived by inserting 3xMYC tag sequence in the downstream of 5’ LTR in pMSCV-puro vector. EGFP-tagged α5-integrin (Laukaitis et al., 2001) was a gift from A. F. Horwitz (University of Virginia, USA). pMSCV-puro EGFP-mPaxillin, previously described (Nader et al., 2016); cDNA for human β1-integrin was obtained by PCR using a template plasmid from Addgene (#69804), the sequence was modified to introduce BamHI and NotI restriction sites, and it was then inserted into a vector using these sites. Human KIF3C cDNA and KIF3C-tail (409-793 aa) constructs were obtained by PCR from HeLa cell mRNA with BamHI and NotI restriction site introduction and were inserted into a vector with these restriction sites. Human RCP cDNA was obtained by PCR from HeLa cell mRNA with BamHI and EcoRI restriction site introduction and were inserted into a vector with these restriction sites. pMSCV-puro EGFP-N4-RCP WT, this manuscript. All constructs were confirmed by DNA sequencing.

Primers which were used to prepare constructs:

5’ primer for inserting a MYC tag cDNA and making a pEF1a-MYC-N4 vector: GGCCGCCGGCGAATTCGAGCAGAAGCTGATCAGCGAGGAGGACCTGT

3’ primer for inserting a MYC tag cDNA and making a pEF1a-MYC-N4 vector: GGCCGCCGGCGAATTCGAGCAGAAGCTGATCAGCGAGGAGGACCTGT

5’ primer for inserting a first half of 3xMYC tag cDNA and making a pMSCV-puro-3xMYC-N4 vector: GGCCGCCGGCGAATTCGAGCAGAAGCTGATCAGCGAGGAGGACCTGGAACA

3’ primer for inserting a first half of 3xMYC tag cDNA and making a pMSCV-puro-3xMYC-N4 vector: CAGCTTCTGTTCCAGGTCCTCCTCGCTGATCAGCTTCTGCTCGAATTCGCCGGC

5’ primer for inserting a second half of 3xMYC tag cDNA and making a pMSCV-puro-3xMYC-N4 vector: GAAGCTGATCTCCGAGGAAGATCTGGAGCAGAAGCTGATTAGCGAAGAGGACCTGT

3’ primer for inserting a second half of 3xMYC tag cDNA and making a pMSCV-puro-3xMYC-N4 vector: AATTACAGGTCCTCTTCGCTAATCAGCTTCTGCTCCAGATCTTCCTCGGAGAT

5’ primer for inserting a BamHI restriction site in 5’ end of hKIF3C WT cDNA: CTTCGGGATCCGCCACCATGGCCAGTAAGACCAAGGCC

5’ primer for inserting a BamHI restriction site in 5’ end of hKIF3C tail cDNA: CTTCGGGATCCGCCACC ATGAGCCGCAGGAAGAAGGCC

3’ primer for inserting a NotI restriction site in 3’ end of hKIF3C WT/tail cDNAs: 5’ primer for inserting a BamHI restriction site in 5’ end of hITGB1 WT cDNA: CCCCCGGATCCGCCACCATGAATTTACAACCAATT

3’ primer for inserting a NotI restriction site in 3’ end of hITGB1 WT cDNA: GATTTGCGGCCGCTTTTCCCTCATACTTCGG

5’ primer for inserting a BamHI restriction site in 5’ end of hRCP WT cDNA: GCGCCGGAATTAGATCTGCT GGATCC GCCACC ATGTCCCTAATGGTCTCG

3’ primer for inserting a EcoRI restriction site in 3’ end of hRCP WT cDNA: CGCTGATCAGCTTCTGCTCGAATTC CATCTTTCCTGCTTTTTTGC

All primers were purchased from Integrated DNA Technologies.

Predesigned shRNAs (MilliporeSigma): Rab11-FIP1 / RCP: TRCN0000028230; TRCN0000028253; TRCN0000028301 KIF3C: TRCN0000116728; TRCN0000116730.

siRNA for Rab11-FIP5 was from MilliporeSigma (EMU054831); siRNA for GFP (negative control) was from MilliporeSigma (EHUEGFP). All other siRNA oligonucleotides were from Shanghai GenePharma Co., Ltd (Shanghai, China). siRNAs were designed using Dharmacon algorithm (http://dharmacon.gelifesciences.com/design-center/). Sequences:

KIF3A: 5’-TGTCATACTTGGAGATATA-3’, 5’-CAGATAAGATGGTGGAAAT-3’ and 5’-GGAAATAACATGAGGAAAC-3’;

KIF3B: 5’-GAGCAGAAACGTCGAGAAA-3’, 5’-GAAGAAGACCATTGGAAAT-3’ and 5’- AGATAGAGCATGTGATGAA-3’;

KIF3C: 5’-CCGAAGAGGAAGATGATAA-3’, 5’-GGCTAGAGCTGAAGGAGAA-3’; 5’- GGAGATTGCGGAACAGAAA-3’.

KAP3A: 5’-GAAGAAAGCTGTTGATGAA-3’, 5’-GAGAAAGGATTATGACAAA-3’ and 5’- AGAGAAGACTGGAAACAAA-3’;

siNC (negative control): 5’-UUCUUCGAACGUGUCACGUTT-3’

### Cell culture

NIH3T3 fibroblasts (ATCC) were cultured in Dulbecco’s Modified Eagle Medium (DMEM, Corning Inc.) with 10 mM HEPES pH 7.4 and 10% (v/v) bovine calf serum (BCS, Corning Inc.) as previously described (Cook et al., 1998; Gundersen et al., 1994). For serum starvation, cells were platted on acid-washed coverslips, grown to confluence (2 days), and transferred to serum-free medium (SFM; DMEM, 10 mM HEPES, pH 7.4) for 2 days as previously described (Cook et al., 1998; Gundersen et al., 1994). HEK 293T cells were cultivated in DMEM with 10 mM HEPES pH 7.4 and 10% (v/v) fetal bovine serum (FBS, Gemini Bio-Products) as previously described (Antoku et al., 2019). Human dermal fibroblasts (HDF) were cultivated in DMEM with 10 mM HEPES pH 7.4 and 15% (v/v) FBS (Gemini Bio-Products) as previously described (Chang et al., 2019). Doxycycline (MilliporeSigma) was added overnight when stated, to induce expression of inducible proteins.

### Microinjection, transfection and viral infection

For microinjection, 10-50 µg/ml purified plasmid DNA in 10 mM HEPES, 140 mM KCl pH 7.4 was pressure microinjected into nuclei of cells at the wounded edge using a glass micropipette and allowed to express for 1 hour. Plasmids were transfected with lipofectamine 3000 following the manufacturer protocol (Thermo Fisher Scientific); siRNAs (20 µM) were transfected with using Lipofectamine RNAiMAX (Thermo Fisher Scientific) according to manufacturer’s instructions for reverse transfection. Viral production and infection were performed as previously described (Antoku et al., 2019)

### Wound healing, FA formation, and FA disassembly and reassembly assays

For the wound healing assay, the cells were serum-starved for 2 days, scratched and allowed to recover for 2 hours before adding BCS (final concentration of 2% BCS (v/v)), followed by fixing and immunofluorescent staining. FA formation was analyzed in 2 day serum-starved NIH3T3 fibroblasts treated with 10 µM LPA (Avantor, Inc.) for 15-90 min. FA disassembly and reassembly after nocodazole washout were assessed as previously described (Ezratty et al., 2005; Nader et al., 2016). Briefly, NIH3T3 fibroblasts were grown on glass coverslips and treated with 10 µM nocodazole (MilliporeSigma) for 3-4 hours to completely depolymerize microtubules. The drug was washed out with SFM, and microtubules allowed to repolymerize for different intervals before fixing and immunofluorescent staining or preparing cell lysates for GST-pulldowns.

### Cell surface labeling and flow cytometry

Cells were harvested by rapid trypsinization at various time points during microtubule regrowth and FA disassembly, and washed in ice-cold PCN (1x PBS, 0.5% BCS, 0.1% NaN_3_). Approximately 4x10^6^ cells per sample were stained without fixation at 4 °C. For detection of α5 integrin in NIH3T3 fibroblasts, PE–conjugated rat α-mouse CD49e monoclonal antibody (BD Biosciences) was used at dilution of 1:1.500. After 3-4 washes to remove unbound antibody, samples were fixed in ice-cold 4% paraformaldehyde and the distribution and mean fluorescence intensity of each sample was quantified using a Becton Dickson FACScan.

### GST-pulldown

Cells were gently washed with ice-cold phosphate-buffered saline (PBS), lysed in lysis buffer [20 mM HEPES, 100 mM NaCl, 20 mM MgCl_2_, and 1% Triton X-100 (pH 7.4)] supplemented with protease and phosphatase inhibitors (Thermo Fisher Scientific), and incubated for 30 minutes on a rocker at 4°C, and the lysates were centrifuged at 15,000 rpm for 15 minutes at 4°C. Supernatants were incubated with GST fusion proteins pre-coupled to Glutathione-Sepharose beads (Cytiva) for 1 hour at 4°C. GST beads attached to the fusion proteins were washed three times with the same lysis buffer, eluted in SDS-PAGE sample buffer, resolved by SDS-PAGE, and processed for immunoblotting.

### Western blotting

Cells lysed in SDS sample buffer were boiled for 5 min and then separated by SDS-PAGE on a either a 10% gel or NuPage 4-12% Bis-Tris gels (Thermo Fisher Scientific). Gels were transferred to nitrocellulose, blocked for 1 hour in 5% non-fat dry milk (LabScientific) and then incubated for 1h-overnight with different primary antibodies followed by 1 hour incubation with the appropriate HRP-conjugated or IRDye-conjugated secondary antibodies. After washing the membrane several times with TBST (50 mM Tris-HCl pH 7.4, 150 mM NaCl, 0.1 % Tween-20 (v/v)), they were scanned and recorded with an Odyssey CLx Imager (LICOR).

### Immunofluorescence

For immunostaining cells were either fixed in 4% paraformaldehyde in PBS for 10 minutes or fixed in -20°C methanol for microtubule staining followed by permeabilization with 0.3% Triton in PBS for 10 minutes and blocking with 2% BSA. Fixed cells were incubated with primary and secondary antibodies (or stained with Alexa Fluor^TM^ 647 phalloidin, Thermo Fisher Scientific). Coverslips were mounted with Fluoromount-G with DAPI (Thermo Fisher Scientific) or PBS when imaging with total internal reflection microscope (TIRFM).

### Fluorescence Microscopy

TIRF and epifluorescence images were acquired with an Apo TIRF 60x oil objective (NA 1.49) and a Nikon DS-Qi2 CMOS camera on a Nikon Eclipse Ti microscope controlled by Nikon’s NIS-Elements software or with an Apo TIRF 60x oil objective (NA 1.49); C-NSTORM QUAD 405/488/561/647 filter set; LUNF 488/561/640 NM 1F TIRF lasers and ORCA-FUSIONBT SCMOS USB3.0 CARD C-MT camera on a Nikon Ti2E Motorized Encoded Stage controlled by Nikon’s NIS-Elements software. Confocal and epifluorescence images were acquired with a Plan Apochromat Lambda D 60x Oil objective (NA 1.45) using Andor (Oxford instruments) BC43 microscope, by Andor Fusion software.

### Image Processing and Quantification

Images were processed and quantified with Image J (NIH, Bethesda, MD); all images were background subtracted before performing measurements. To analyze FA size and number a ROI was set around each cell and a fixed threshold was set to all the images in the same experiment, followed by analyze particles (size=0.40-15.00; circularity=0.00-0.99). To analyze the distribution of adhesions we used analyze > plot profile, a ROI was set around cells that had its sides surrounded by other cells and had the front of the cell at ∼90° from the wound. Integrin and KIF3A in the Rab11 ERC after FA disassembly, were quantified as previously described (Nader et al., 2016). Live imaging of cells expressing RCP-EGFP (Supplementary Movie 3) were processed with “Denoise.AI” from Nikon NIS-Elements, followed by “equalize intensity in time”, to correct for photobleaching. The videos were converted to maximum projection using Nikon NIS-Elements. Clear vesicle paths longer than 2 µm were manually counted. Cells within monolayer (away from the wound edge) were randomly divided into two equal halves, whereas cells at the wound edge were divided into two equal halves parallel to the wound edge (representing front and back of the cell).

### Statistical analysis

A two-tailed unpaired Student’s *t* test was used to evaluate two groups. Significance between multiple groups was determined by one-way analysis of variance (ANOVA) followed by Tukey’s multiple comparison test, using GraphPad Prism 10. Error bars, mean ± SD; p values: *, < 0.05; **, < 0.01; ***, <0.005; ****, 0.001; NS, not significant.

## Supplementary material

Our findings indicate that KIF3A and KIF3C are not essential for the initial formation of FAs. Interestingly, the KIF3A adaptors, KAP3 and Rab11-FIP5 were found to be not required for FA reassembly after nocodazole washout. In addition to these findings, this section also includes Western blot demonstrating the knockdown of proteins used throughout the study. These results provide crucial validation of our experimental approach and further support our conclusions.

## Acknowledgements

We thank Dr. Geri Kreitzer (CUNY, New York) for the KIF3A-tail-EGFP construct. Research reported in this publication was supported by NIH grant R35 GM136403 to G.G.G. The content is solely the responsibility of the authors and does not necessarily represent the official views of the NIH.

## Abbreviations

FA: Focal adhesion
ERC: Endocytic recycling compartment
NZWO: Nocodazole washout
LPA: lysophosphatidic acid
HDF: human dermal fibroblasts.

## Supplementary figures

**Supplementary figure 1: The Kinesin-2 family member KIF3A and KIF3C is not required for FA formation. A)** Western blot of KIF3A in NIH3T3 fibroblasts treated with noncoding siRNA (siNC) or two different KIF3A siRNAs (siKIF3A.1 or siKIF3A.2). Tubulin was used as a loading control. Molecular weight markers in kDa are on the right (also in panels B, C). **B)** Western blot of KIF3B in NIH3T3 fibroblasts treated with the indicated siRNAs. Tubulin was used as a loading control. **C)** Western blot of KIF3C in NIH3T3 fibroblasts treated with the indicated siRNAs. Tubulin is a loading control. **D)** Representative images of FA formation in serum-starved NIH3T3 fibroblast monolayers wounded and stimulated with 10 µM LPA for 15 or 90 min. Cells were treated with siRNAs and stained for paxillin and filamentous actin as in A. **E)** Quantification of 90 cells per condition of FA number and size for cells treated as in D. The histogram bars represent the SD, with each dot indicating the mean of an individual experiment. **F)** Representative images of microtubules in NIH3T3 fibroblasts at 0, 60 or 120 min after nocodazole washout. Cells were treated with the indicated siRNAs and stained with tubulin antibody. Scale bar, 10µm. **G)** Representative images of vinculin-stained FAs in the same cells depicted in F. Scale bar, 10µm. **H)** Quantification of cells with more than 10 FAs from cells treated as in G. At least 150 cells were quantified per condition from 5 independent experiments.

**Supplementary Figure 2: The Kinesin-2 family member KIF3A and KIF3C are required for FA reassembly after nocodazole washout. A)** Western blots of human dermal fibroblasts transfected with the indicated shRNAs. Western blots were stained with the antibodies on the left. Tubulin was used as a loading control. **B)** Representative images of human dermal fibroblasts from a nocodazole washout assay to assess microtubule-induced FA disassembly and reassembly. Cells fixed at 0, 45 and 120 min after nocodazole washout and stained with paxillin antibody. Cells were transfected with the indicated shRNAs before the nocodazole washout assay. Scale bar, 15 µm. **C)** Quantification of paxillin-stained FA number and size in human dermal fibroblasts as shown in B. Data are from 30 cells per time point from 3 independent experiments. The histogram bars represent the SD, with each dot indicating the mean of an individual experiment.

**Supplementary Figure 3: KIF3A adaptors KAP3 and Rab11-FIP5 are not required for FA reassembly A)** Representative images of paxillin-stained FAs in NIH3T3 fibroblasts at the indicated times after nocodazole washout. Before the nocodazole assay, cells were treated with siRNAs to GFP (as control) or Rab11-FIP5. Scale bar, 15µm. **B)** Quantification of paxillin-stained FA number and size for the cells treated as in A (time is min after nocodazole washout). Data are from 30 cells per time point from 3 independent experiments. The histogram bars represent the SD, with each dot indicating the mean of an individual experiment. **C)** Western blots of NIH3T3 fibroblasts treated with the indicated siRNAs. Western blots were stained with the antibodies indicated on the left; molecular weight markers are on the right. Tubulin is a loading control. **D)** Representative images of vinculin-stained FAs and microtubules in NIH3T3 fibroblasts at the indicated times after nocodazole washout. Before nocodazole washout, cells were treated with the indicated siRNAs. Scale bar, 10µm. **E)** Quantification of cells with >10 FAs for cells treated as depicted in D. Data are from at least 150 cells per time point from 3 independent experiments. The histogram bars represent the SD, with each dot indicating the mean of an individual experiment. **F, G)** Western blots of NIH3T3 fibroblasts treated with the indicated siRNAs or shRNAs. Western blots were stained with the antibodies indicated on the left. Molecular weight markers are on the right. Tubulin is a loading control. shRCP.2 was excluded from experiments because it exhibited insufficient knockdown efficiency. Scale bar 10µm.

**Supplementary Movie 1:** Focal adhesion behavior during nocodazole washout in NIH3T3 fibroblasts treated with siNC or siKIF3C and expressing Paxillin-GFP. Images were acquired by TIRF microscopy every 1 min. Time in min. Scale bar 10µm.

**Supplementary Movie 2:** Focal adhesion behavior during nocodazole washout in NIH3T3 fibroblasts treated with shNC or shRCP and expressing paxillin-GFP. Images were acquired by TIRF microscopy every 1 min. Time in min. Scale bar 10µm.

**Supplementary Movie 3:** Movies of NIH3T3 fibroblasts expressing RCP-EGFP after nocodazole washout used to generate maximum projections of RCP vesicle movements in Figure 6F. Movies show polarized cells at the wound edge and nonpolarized cells within the monolayer. Images were acquired by HILO microscopy every 400 msec. Time in sec. Scale bar 10µm.

## References

Antoku, S., Wu, W., Joseph, L.C., Morrow, J.P., Worman, H.J., Gundersen, G.G., 2019. ERK1/2 phosphorylation of FHOD connects signaling and nuclear positioning alternations in cardiac laminopathy. Dev. Cell 51, 602–616.e12. 10.1016/j.devcel.2019.10.023

Boehlke, C., Kotsis, F., Buchholz, B., Powelske, C., Eckardt, K.-U., Walz, G., Nitschke, R., Kuehn, E.W., 2013. Kif3a Guides Microtubular Dynamics, Migration and Lumen Formation of MDCK Cells. PLoS ONE 8, e62165. 10.1371/journal.pone.0062165

Bridgewater, R.E., Norman, J.C., Caswell, P.T., 2012. Integrin trafficking at a glance. J. Cell Sci. 125, 3695– 3701. 10.1242/jcs.095810

Caswell, P., Norman, J., 2008. Endocytic transport of integrins during cell migration and invasion. Trends Cell Biol. 18, 257–263. 10.1016/j.tcb.2008.03.004

Caswell, P.T., Chan, M., Lindsay, A.J., McCaffrey, M.W., Boettiger, D., Norman, J.C., 2008. Rab-coupling protein coordinates recycling of α5β1 integrin and EGFR1 to promote cell migration in 3D microenvironments. J. Cell Biol. 183, 143–155. 10.1083/jcb.200804140

Chang, W., Wang, Y., Luxton, G.W.G., Östlund, C., Worman, H.J., Gundersen, G.G., 2019. Imbalanced nucleocytoskeletal connections create common polarity defects in progeria and physiological aging. Proc. Natl. Acad. Sci. 116, 3578–3583. 10.1073/pnas.1809683116

Chao, W.-T., Kunz, J., 2009. Focal adhesion disassembly requires clathrin-dependent endocytosis of integrins. FEBS Lett. 583, 1337–1343. 10.1016/j.febslet.2009.03.037

Conway, J.R.W., Jacquemet, G., 2019. Cell matrix adhesion in cell migration. Essays Biochem. 63, 535–551. 10.1042/EBC20190012

De Franceschi, N., Hamidi, H., Alanko, J., Sahgal, P., Ivaska, J., 2015. Integrin traffic – the update. J. Cell Sci. 128, 839–852. 10.1242/jcs.161653

Doyle, A.D., Wang, F.W., Matsumoto, K., Yamada, K.M., 2009. One-dimensional topography underlies three-dimensional fibrillar cell migration. J. Cell Biol. 184, 481–490. 10.1083/jcb.200810041

Eskova, A., Knapp, B., Matelska, D., Reusing, S., Arjonen, A., Lisauskas, T., Pepperkok, R., Russell, R., Eils, R., Ivaska, J., Kaderali, L., Erfle, H., Starkuviene, V., 2014. An RNAi screen identifies KIF15 as a novel regulator of the endocytic trafficking of integrin. J. Cell Sci. 127, 2433–2447. 10.1242/jcs.137281

Eva, R., Dassie, E., Caswell, P.T., Dick, G., ffrench-Constant, C., Norman, J.C., Fawcett, J.W., 2010. Rab11 and Its Effector Rab Coupling Protein Contribute to the Trafficking of β1 Integrins during Axon Growth in Adult Dorsal Root Ganglion Neurons and PC12 Cells. J. Neurosci. 30, 11654–11669. 10.1523/JNEUROSCI.2425-10.2010

Ezratty, E.J., Bertaux, C., Marcantonio, E.E., Gundersen, G.G., 2009. Clathrin mediates integrin endocytosis for focal adhesion disassembly in migrating cells. J. Cell Biol. 187, 733–747. 10.1083/jcb.200904054

Ezratty, E.J., Partridge, M.A., Gundersen, G.G., 2005. Microtubule-induced focal adhesion disassembly is mediated by dynamin and focal adhesion kinase. Nat. Cell Biol. 7, 581–590. 10.1038/ncb1262

Fanfone, D., Wu, Z., Mammi, J., Berthenet, K., Neves, D., Weber, K., Halaburkova, A., Virard, F., Bunel, F., Jamard, C., Hernandez-Vargas, H., Tait, S.W., Hennino, A., Ichim, G., 2022. Confined migration promotes cancer metastasis through resistance to anoikis and increased invasiveness. eLife 11, e73150. 10.7554/eLife.73150

Ferro, E., Bosia, C., Campa, C.C., 2021. RAB11-Mediated Trafficking and Human Cancers: An Updated Review. Biology 10, 26. 10.3390/biology10010026

Gao, Y., Zheng, H., Li, L., Zhou, C., Chen, X., Zhou, X., Cao, Y., 2020. KIF3C Promotes Proliferation, Migration, and Invasion of Glioma Cells by Activating the PI3K/AKT Pathway and Inducing EMT. BioMed Res. Int. 2020, 6349312. 10.1155/2020/6349312

Garbouchian, A., Montgomery, A.C., Gilbert, S.P., Bentley, M., 2022. KAP is the neuronal organelle adaptor for Kinesin-2 KIF3AB and KIF3AC. Mol. Biol. Cell 33, ar133. 10.1091/mbc.E22-08-0336

Gardel, M.L., Schneider, I.C., Yvonne Aratyn-Schaus, Waterman, C.M., 2010. Mechanical Integration of Actin and Adhesion Dynamics in Cell Migration. Annu. Rev. Cell Dev. Biol. 26, 315–333. 10.1146/annurev.cellbio.011209.122036

Gilbert, S.P., Guzik-Lendrum, S., Rayment, I., 2018. Kinesin-2 motors: Kinetics and biophysics. J. Biol. Chem. 293, 4510–4518. 10.1074/jbc.R117.001324

Gundersen, G.G., Bulinski, J.C., 1988. Selective stabilization of microtubules oriented toward the direction of cell migration. Proc. Natl. Acad. Sci. U. S. A. 85, 5946–5950.

Gundersen, G.G., Kalnoski, M.H., Bulinski, J.C., 1984. Distinct populations of microtubules: tyrosinated and nontyrosinated alpha tubulin are distributed differently in vivo. Cell 38, 779–789. 10.1016/0092-8674(84)90273-3

Hammond, J.W., Blasius, T.L., Soppina, V., Cai, D., Verhey, K.J., 2010. Autoinhibition of the kinesin-2 motor KIF17 via dual intramolecular mechanisms. J. Cell Biol. 189, 1013–1025. 10.1083/jcb.201001057

He, Z., Wang, J., Xu, J., Jiang, X., Liu, X., Jiang, J., 2022. Dynamic regulation of KIF15 phosphorylation and acetylation promotes focal adhesions disassembly in pancreatic cancer. Cell Death Dis. 13, 1–13. 10.1038/s41419-022-05338-y

Hirokawa, N., Noda, Y., Tanaka, Y., Niwa, S., 2009. Kinesin superfamily motor proteins and intracellular transport. Nat. Rev. Mol. Cell Biol. 10, 682–696. 10.1038/nrm2774

Huet-Calderwood, C., Rivera-Molina, F., Iwamoto, D.V., Kromann, E.B., Toomre, D., Calderwood, D.A., 2017. Novel ecto-tagged integrins reveal their trafficking in live cells. Nat. Commun. 8, 570. 10.1038/s41467-017-00646-w

Lauffenburger, D.A., Horwitz, A.F., 1996. Cell Migration: A Physically Integrated Molecular Process. Cell 84, 359–369. 10.1016/S0092-8674(00)81280-5

Laukaitis, C.M., Webb, D.J., Donais, K., Horwitz, A.F., 2001. Differential dynamics of alpha 5 integrin, paxillin, and alpha-actinin during formation and disassembly of adhesions in migrating cells. J. Cell Biol. 153, 1427–1440. 10.1083/jcb.153.7.1427

Li, D., Kuehn, E.W., Prekeris, R., 2014. Kinesin-2 mediates apical endosome transport during epithelial lumen formation. Cell. Logist. 4, e28928. 10.4161/cl.28928

Li, R., Gundersen, G.G., 2008. Beyond polymer polarity: how the cytoskeleton builds a polarized cell. Nat. Rev. Mol. Cell Biol. 9, 860–873. 10.1038/nrm2522

Liao, G., Gundersen, G.G., 1998. Kinesin is a candidate for cross-bridging microtubules and intermediate filaments. Selective binding of kinesin to detyrosinated tubulin and vimentin. J. Biol. Chem. 273, 9797– 9803. 10.1074/jbc.273.16.9797

Lin, S.X., Gundersen, G.G., Maxfield, F.R., 2002. Export from pericentriolar endocytic recycling compartment to cell surface depends on stable, detyrosinated (glu) microtubules and kinesin. Mol. Biol. Cell 13, 96–109. 10.1091/mbc.01-05-0224

Luster, A.D., Alon, R., von Andrian, U.H., 2005. Immune cell migration in inflammation: present and future therapeutic targets. Nat. Immunol. 6, 1182–1190. 10.1038/ni1275

Marszalek, J.R., Goldstein, L.S.B., 2000. Understanding the functions of kinesin-II. Biochim. Biophys. Acta BBA - Mol. Cell Res. 1496, 142–150. 10.1016/S0167-4889(00)00015-X

McKenney, R.J., Huynh, W., Vale, R.D., Sirajuddin, M., 2016. Tyrosination of α-tubulin controls the initiation of processive dynein–dynactin motility. EMBO J. 35, 1175–1185. 10.15252/embj.201593071

Mitchison, T.J., Cramer, L.P., 1996. Actin-based cell motility and cell locomotion. Cell 84, 371–379. 10.1016/s0092-8674(00)81281-7

Möhl, C., Kirchgessner, N., Schäfer, C., Hoffmann, B., Merkel, R., 2012. Quantitative mapping of averaged focal adhesion dynamics in migrating cells by shape normalization. J. Cell Sci. 125, 155–165. 10.1242/jcs.090746

Moon, H.H., Kreis, N.-N., Friemel, A., Roth, S., Schulte, D., Solbach, C., Louwen, F., Yuan, J., Ritter, A., 2021. Mitotic Centromere-Associated Kinesin (MCAK/KIF2C) Regulates Cell Migration and Invasion by Modulating Microtubule Dynamics and Focal Adhesion Turnover. Cancers 13, 5673. 10.3390/cancers13225673

Moreno-Layseca, P., Icha, J., Hamidi, H., Ivaska, J., 2019. Integrin trafficking in cells and tissues. Nat. Cell Biol. 21, 122–132. 10.1038/s41556-018-0223-z

Muresan, V., Abramson, T., Lyass, A., Winter, D., Porro, E., Hong, F., Chamberlin, N.L., Schnapp, B.J., 1998. KIF3C and KIF3A form a novel neuronal heteromeric kinesin that associates with membrane vesicles. Mol. Biol. Cell 9, 637–652. 10.1091/mbc.9.3.637

Nader, G.P.F., Ezratty, E.J., Gundersen, G.G., 2016. FAK, talin and PIPKIγ regulate endocytosed integrin activation to polarize focal adhesion assembly. Nat. Cell Biol. 18, 491–503. 10.1038/ncb3333

Pawluchin, A., Galic, M., 2022. Moving through a changing world: Single cell migration in 2D vs. 3D. Front. Cell Dev. Biol. 10, 1080995. 10.3389/fcell.2022.1080995

Petrie, R.J., Doyle, A.D., Yamada, K.M., 2009. Random versus directionally persistent cell migration. Nat. Rev. Mol. Cell Biol. 10, 538–549. 10.1038/nrm2729

Regen, C.M., Horwitz, A.E., 1992. Dynamics of/ 1 Integrin-mediated Adhesive Contacts in Motile Fibroblasts. J. Cell Biol. 119.

Ridley, A.J., Hall, A., 1992. The small GTP-binding protein rho regulates the assembly of focal adhesions and actin stress fibers in response to growth factors. Cell 70, 389–399. 10.1016/0092-8674(92)90163-7

Ridley, A.J., Schwartz, M.A., Burridge, K., Firtel, R.A., Ginsberg, M.H., Borisy, G., Parsons, J.T., Horwitz, A.R., 2003. Cell migration: integrating signals from front to back. Science 302, 1704–1709. 10.1126/science.1092053

Scarpa, E., Mayor, R., 2016. Collective cell migration in development. J. Cell Biol. 212, 143–155. 10.1083/jcb.201508047

Schonteich, E., Wilson, G.M., Burden, J., Hopkins, C.R., Anderson, K., Goldenring, J.R., Prekeris, R., 2008a. The Rip11/Rab11-FIP5 and kinesin II complex regulates endocytic protein recycling. J. Cell Sci. 121, 3824– 3833. 10.1242/jcs.032441

Schonteich, E., Wilson, G.M., Burden, J., Hopkins, C.R., Anderson, K., Goldenring, J.R., Prekeris, R., 2008b. The Rip11/Rab11-FIP5 and kinesin II complex regulates endocytic protein recycling. J. Cell Sci. 121, 3824– 3833. 10.1242/jcs.032441

Sirajuddin, M., Rice, L.M., Vale, R.D., 2014. Regulation of microtubule motors by tubulin isotypes and post-translational modifications. Nat. Cell Biol. 16, 335–344. 10.1038/ncb2920

Smilenov, L.B., Mikhailov, A., Pelham, R.J., Marcantonio, E.E., Gundersen, G.G., 1999. Focal adhesion motility revealed in stationary fibroblasts. Science 286, 1172–1174. 10.1126/science.286.5442.1172

Stehbens, S.J., Paszek, M., Pemble, H., Ettinger, A., Gierke, S., Wittmann, T., 2014. CLASPs link focal-adhesion-associated microtubule capture to localized exocytosis and adhesion site turnover. Nat. Cell Biol. 16, 558–570. 10.1038/ncb2975

Theisen, U., Straube, E., Straube, A., 2012. Directional persistence of migrating cells requires Kif1C- mediated stabilization of trailing adhesions. Dev. Cell 23, 1153–1166. 10.1016/j.devcel.2012.11.005

Vicente-Manzanares, M., Ma, X., Adelstein, R.S., Horwitz, A.R., 2009. Non-muscle myosin II takes centre stage in cell adhesion and migration. Nat. Rev. Mol. Cell Biol. 10, 778–790. 10.1038/nrm2786

Webster, D.R., Gundersen, G.G., Bulinski, J.C., Borisy, G.G., 1987. Assembly and turnover of detyrosinated tubulin in vivo. J. Cell Biol. 105, 265–276. 10.1083/jcb.105.1.265

Wilcke, M., Johannes, L., Galli, T., Mayau, V., Goud, B., Salamero, J., 2000. Rab11 Regulates the Compartmentalization of Early Endosomes Required for Efficient Transport from Early Endosomes to the Trans-Golgi Network. J. Cell Biol. 151, 1207–1220.

Wilson, B., Flett, C., Gemperle, J., Lawless, C., Hartshorn, M., Hinde, E., Harrison, T., Chastney, M., Taylor, S., Allen, J., Norman, J.C., Zacharchenko, T., Caswell, P.T., 2023. Proximity labelling identifies pro-migratory endocytic recycling cargo and machinery of the Rab4 and Rab11 families. J. Cell Sci. 136, jcs260468. 10.1242/jcs.260468

Yamada, K.M., Sixt, M., 2019. Mechanisms of 3D cell migration. Nat. Rev. Mol. Cell Biol. 20, 738–752. 10.1038/s41580-019-0172-9

Yamazaki, H., Nakata, T., Okada, Y., Hirokawa, N., 1995. KIF3A/B: a heterodimeric kinesin superfamily protein that works as a microtubule plus end-directed motor for membrane organelle transport. J. Cell Biol. 130, 1387–1399. 10.1083/jcb.130.6.1387

Yeo, M.G., Partridge, M.A., Ezratty, E.J., Shen, Q., Gundersen, G.G., Marcantonio, E.E., 2006. Src SH2 Arginine 175 Is Required for Cell Motility: Specific Focal Adhesion Kinase Targeting and Focal Adhesion Assembly Function. Mol. Cell. Biol. 26, 4399–4409. 10.1128/MCB.01147-05

