## Supplementary figures and images for "The kinesin KIF3AC recycles endocytosed integrin to polarize adhesion formation towards the leading edge"

### Supplementary Figure 1

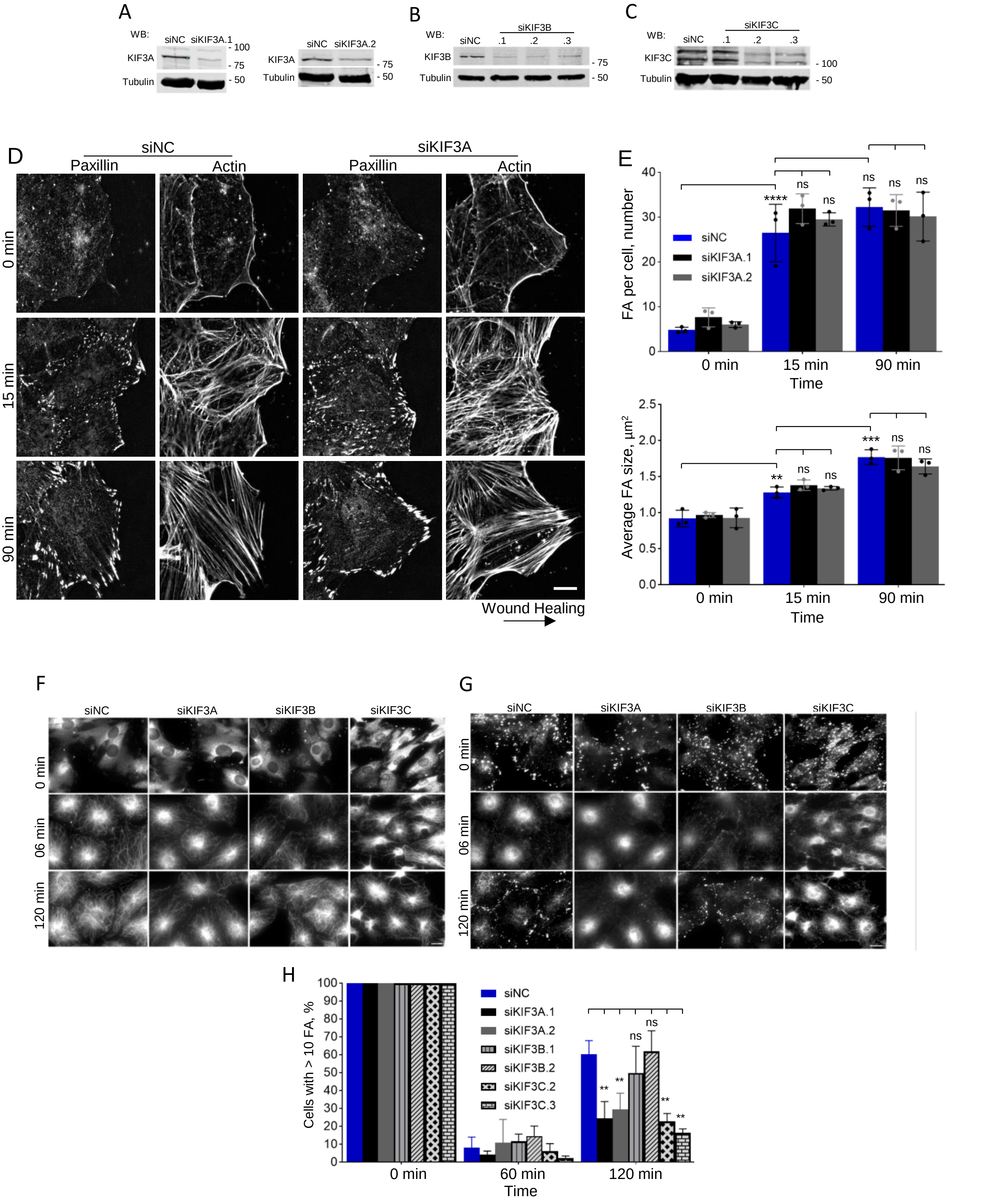

### Supplementary Figure 2

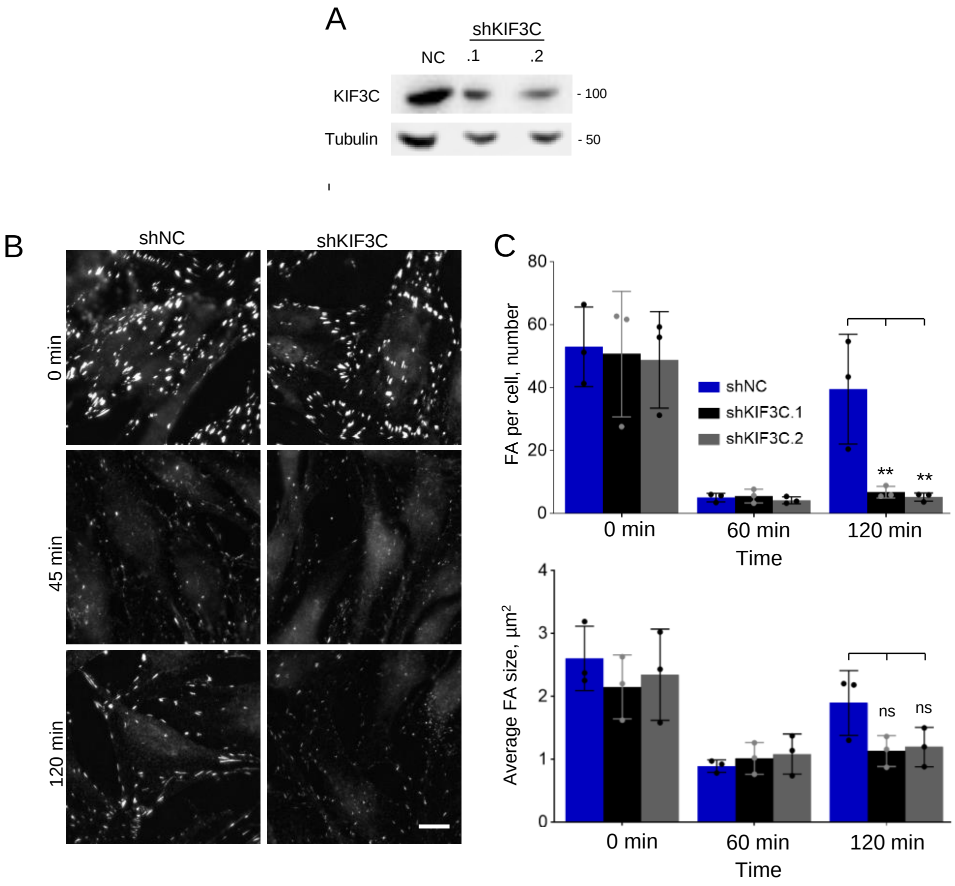

### Supplementary Figure 3

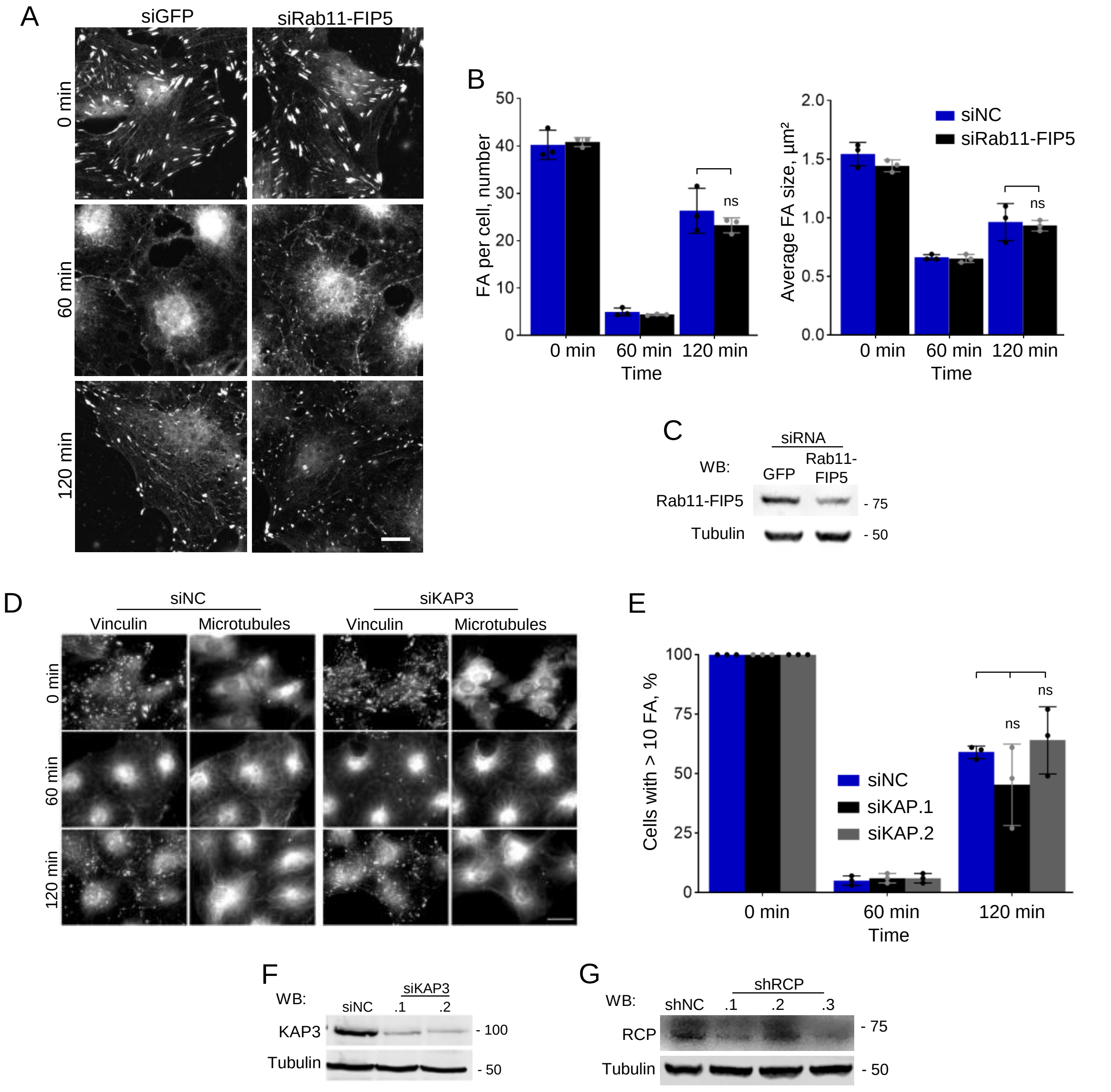
